# CTCF is a DNA-tension-dependent barrier to cohesin-mediated DNA loop extrusion

**DOI:** 10.1101/2022.09.08.507093

**Authors:** Iain F. Davidson, Roman Barth, Maciej Zaczek, Jaco van der Torre, Wen Tang, Kota Nagasaka, Richard Janissen, Jacob Kerssemakers, Gordana Wutz, Cees Dekker, Jan-Michael Peters

**Affiliations:** Research Institute of Molecular Pathology; Vienna BioCenter, 1030 Vienna, Austria; Department of Bionanoscience, Kavli Institute of Nanoscience Delft, Delft University of Technology; Delft, Netherlands

**Author notes:** Children’s Cancer Research Institute, St. Anna Kinderkrebsforschung; 1090 Vienna, Austria. These authors contributed equally to this work (Iain F. Davidson, Roman Barth, Maciej Zaczek).

## Abstract

In eukaryotes, genomic DNA is extruded into loops by cohesin^1^. By restraining this process, the DNA-binding protein CTCF generates topologically associating domains (TADs)^2-4^ that play key roles in gene regulation and recombination during development and disease^1,5-8^. How CTCF establishes TAD boundaries and to what extent these are permeable to cohesin is unknown^9^. To address these questions, we visualize interactions of single CTCF and cohesin molecules on DNA in vitro. We show that CTCF is sufficient to block diffusing cohesin, possibly reflecting how cohesive cohesin accumulates at TAD boundaries, as well as to block loop-extruding cohesin, reflecting how CTCF establishes TAD boundaries. CTCF functions asymmetrically, as predicted, but unexpectedly is dependent on DNA tension. Moreover, CTCF regulates cohesin’s loop extrusion activity by changing its direction and by inducing loop shrinkage. Our data indicate that CTCF is not, as previously assumed, simply a barrier to cohesin-mediated loop extrusion but is an active regulator of this process, where the permeability of TAD boundaries can be modulated by DNA tension. These results reveal mechanistic principles of how CTCF controls loop extrusion and genome architecture.

## Introduction

The folding of genomic DNA by cohesin has important roles in chromatin organization, gene regulation, and recombination ^1^. Cohesin belongs to the ‘structural maintenance of chromosomes’ (SMC) family of ATPase complexes that can extrude DNA into loops, an activity which has been reconstituted *in vitro* for cohesin, condensin, and SMC5/6 ^10-15^. Cohesin also performs a second function by mediating sister chromatid cohesion. In individual cells, loops are located at variable positions, suggesting that loops are dynamic structures most of which are in the process of being extruded ^16-18^. However, in cell-population measurements patterns emerge which reveal that most loops are formed within topologically associating domains (TADs) ^17,19,20^. CTCF (CCCTC binding factor) is located at TAD boundaries ^19,20^ and is required for their formation and for cohesin accumulation at these sites^2-4^. CTCF has unstructured N- and C-terminal regions that flank eleven zinc fingers, several of which recognize an asymmetric DNA sequence and thus position CTCF directionally on DNA ^21,22^. Most CTCF binding sites are oriented in convergent orientations so that CTCF’s N-termini face the interior of TADs, suggesting that CTCF acts as an asymmetric boundary to cohesin-mediated loop extrusion ^23-25^. Consistent with this possibility, CTCF’s N-terminus can bind cohesin ^26^ and is required for TAD insulation and loop anchoring at these sites ^26-29^.

Several mechanisms have been suggested for how CTCF might prevent loop extrusion across TAD boundaries (reviewed in ^9^), namely, as a physical barrier (‘roadblock’), by binding cohesin, by preventing release of cohesin from DNA, by promoting replacement of cohesin’s ATPase activating subunit NIPBL by its inactive counterpart PDS5, by directly inhibiting cohesin’s ATPase activity, and by promoting entrapment of DNA inside a ring structure that is formed by three of cohesin’s subunits ^30^. It has also been proposed that CTCF converts cohesin into an asymmetrically extruding enzyme by stalling loop extrusion on the CTCF-bound site while allowing cohesin to continue reeling DNA into the loop only from the TAD interior ^26,31,32^. However, it remains unresolved which of these proposed mechanisms is used by CTCF and whether CTCF is sufficient for blocking loop extrusion by cohesin. Answering these questions is of great importance since CTCF is required for controlling enhancer-promoter interactions ^1^, cell differentiation, nuclear reprogramming ^7^, recombination of antigen receptor genes ^5,6^, and timing of DNA replication ^33^, and because CTCF mutations have been implicated in tumorigenesis ^8^. CTCF boundaries are also sites at which replicated DNA molecules are connected by cohesin complexes which mediate cohesion ^34^.

### CTCF finds its DNA binding site by facilitated diffusion

To obtain insight into how CTCF controls cohesin, we developed *in vitro* assays in which CTCF-cohesin interactions can be visualized on DNA at the single-molecule level in real time. We first analyzed how CTCF finds its DNA consensus sequence. Consistent with previous reports ^35,36^, recombinant human CTCF (Fig. 1a) bound specifically to DNA oligonucleotides containing a single CTCF binding site in electrophoretic mobility shift assays (EMSAs) in a manner that was reduced by DNA methylation (Fig. 1b). We introduced this CTCF binding site into linear 26.1 kb DNA molecules, tethered these at both ends to glass surfaces in flow cells, stained with Sytox Green, and imaged the DNA molecules using Highly Inclined and Laminated Optical sheet (HILO) microscopy. Following introduction of fluorophore-labeled CTCF, both immobile and mobile CTCF foci were observed at various positions along the DNA (Fig. 1c, d and Extended Data Fig. 1a). CTCF foci at the CTCF binding site were detectable for much longer than those elsewhere, where CTCF proteins often dissociated rapidly unless they arrived at the CTCF binding site while diffusing along DNA (Fig. 1c, d). Most CTCF foci that were not located at the CTCF binding site were removed by a brief salt wash, unlike those at the CTCF binding site (Extended Data Fig. 1b). Fluorescence intensity and photobleaching analysis indicated that these remaining CTCF molecules were monomers (Extended Data Fig. 1c, d). Once bound to their binding sites, the mean residence time of CTCF molecules was ∼32 min (Extended Data Fig. 1e, f). Since most studies reported shorter chromatin residence times in cells ^37-42^ (for an exception see ^40^) we also performed inverse fluorescence recovery after photobleaching (iFRAP) in HeLa cells in which all CTCF alleles had been modified to express GFP-tagged CTCF. We identified dynamically (∼7 min) and stably bound (∼95 min) populations (Extended Data Fig. 2). These results indicate that CTCF finds its DNA binding site by facilitated diffusion and associates with it *in vitro* and *in vivo* over various, but in some cases long, timescales, which raises the possibility that regulation of CTCF’s residence time could contribute to the permeability of TAD boundaries.

**Figure 1.**
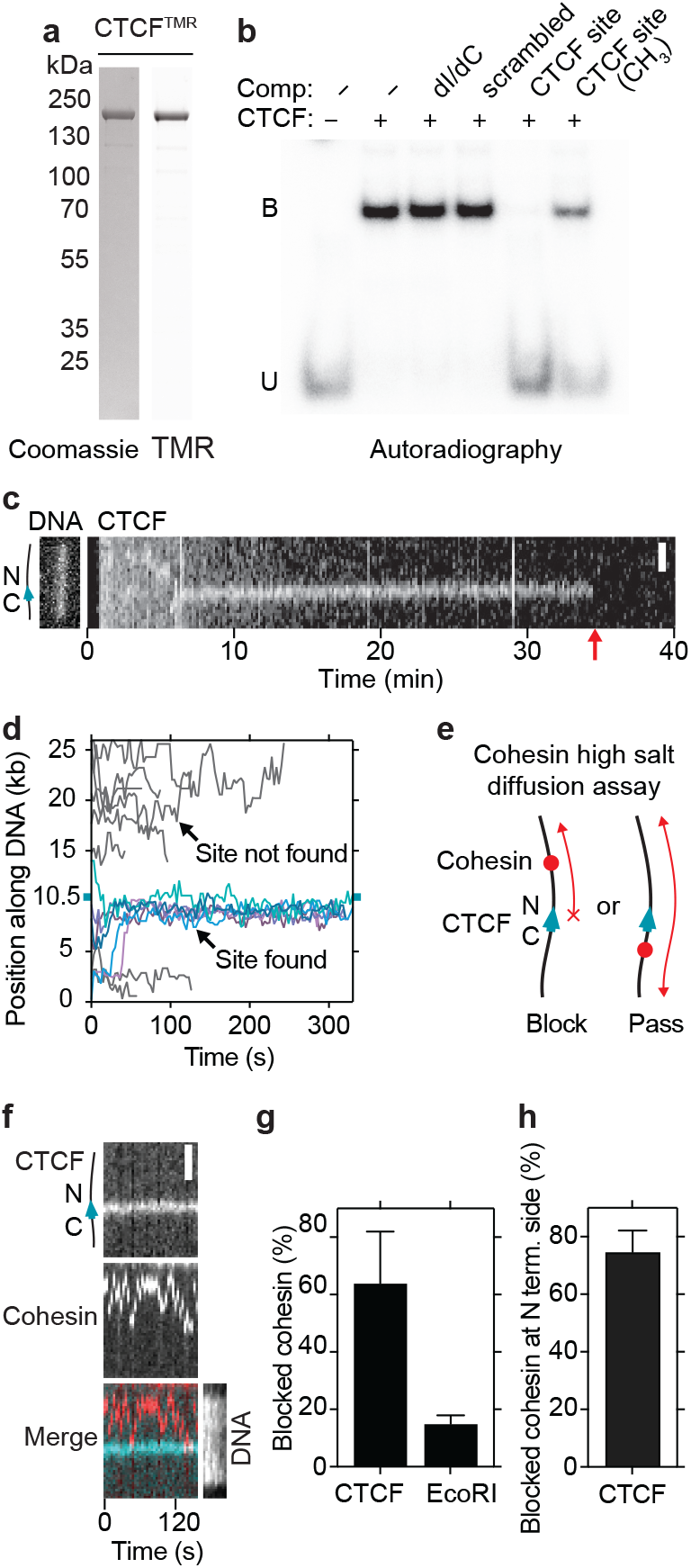
CTCF is a directional barrier to cohesin diffusion on DNA. **a**, Coomassie staining of recombinant CTCF after SDS-PAGE. Tetramethylrhodamine (TMR) was visualized by epi-green excitation. **b**, Autoradiograph of Electrophoretic mobility shift competition assay (EMSA). CTCF was incubated with a ^32^P labelled 100 bp DNA containing a high affinity CTCF binding site. Where indicated, reactions were supplemented with a 100-fold excess of unlabelled DNA competitors. dI-dC: poly(2′-deoxyinosinic-2′-deoxycytidylic acid). B: bound, U: unbound. **c**, Example of TMR-labelled CTCF molecules diffusing on DNA. Non-specifically bound CTCF molecules exhibit random diffusion and dissociate rapidly. At t = ∼ 5.5 min, a CTCF molecule binds DNA and diffuses until it encounters the CTCF binding site at t = 6 min. Scale bar, 2 μm. Red arrow indicates timepoint when CTCF bleached or dissociated. **d**, Superposition of individually tracked TMR-labelled CTCF-diffusion events. Events in which CTCF localized to its binding site at position 10,452 bp (cyan tick) are shown in blue (N = 6). DNA-binding events in which CTCF failed to localize to its binding site are shown in grey (N = 11). **e**, Illustration of cohesin passive diffusion assay. **f**, Example of cohesin diffusion that is blocked by CTCF. Cohesin and CTCF were labeled with Alexa660 (red) and TMR (blue), respectively. Sytox Green DNA stain was introduced into the flow cell at the end of the experiment. Scale bar, 2 μm. **g**, Fraction of blocking events observed when cohesin encountered CTCF or EcoRI^E111Q^ (mean ± SD from 7 (N=264) and 3 (N=106) independent experiments per condition, respectively). **h**, Fraction of blocked events where cohesin diffused on the DNA between the tether point and the N-terminal side of CTCF (mean ± SD from 3 (N=48) independent experiments). In the remaining 25 % of events, cohesin diffused between the tether point and the C-terminal side of CTCF.

**Figure 2.**
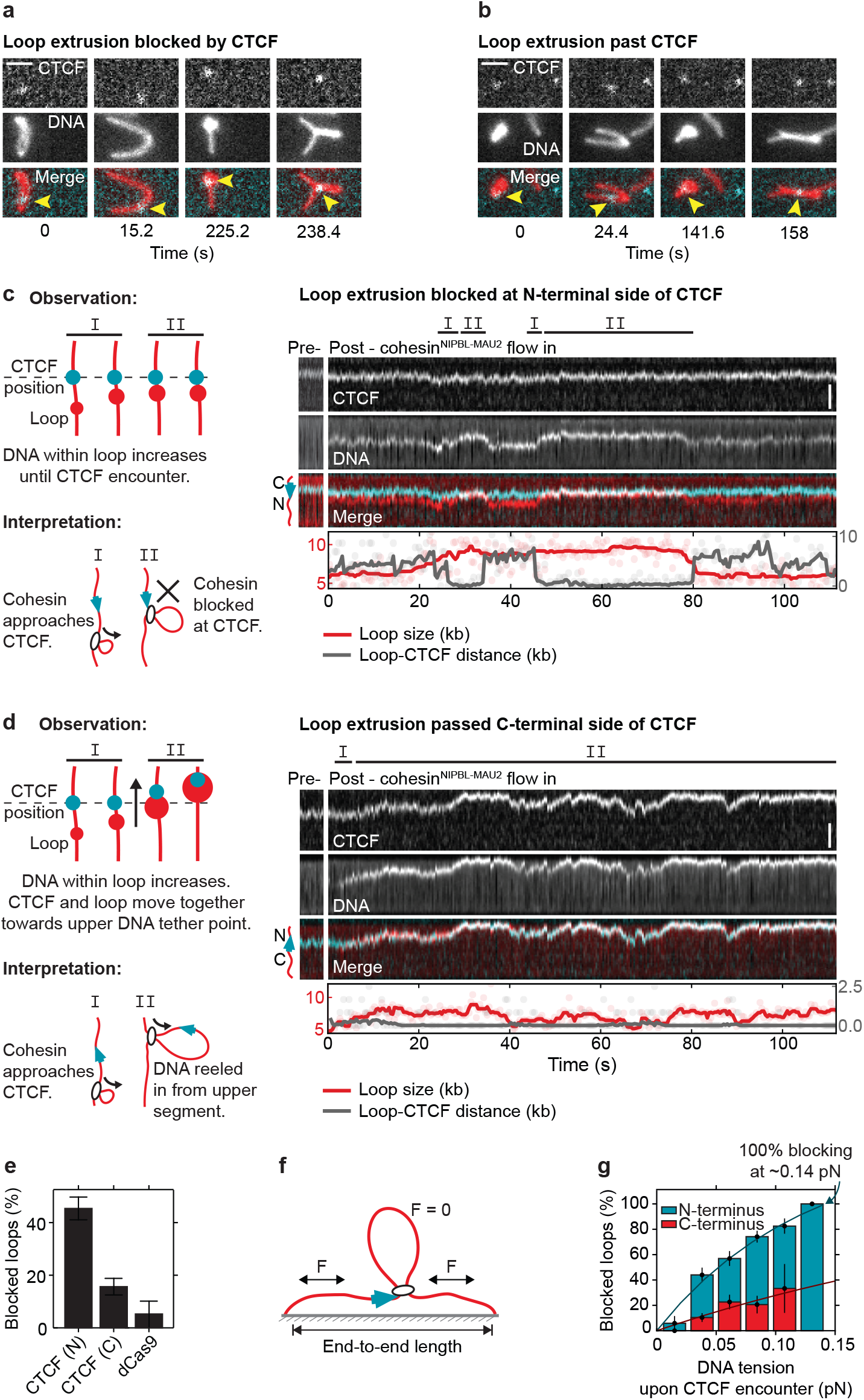
CTCF is a direction- and tension-dependent barrier to cohesin-mediated DNA loop extrusion. **a, b**, Examples of loop extrusion blocked by (a) or passing (b) CTCF (cyan) labeled with Janelia Fluor 646 (JF646). DNA loops (red) were visualized by Sytox Orange stain and buffer flow perpendicular to the DNA axis. Scale bar, 2 μm. **c**, (Left panel) observation and interpretation illustrations of (right panel) a kymograph of cohesin-mediated DNA loop extrusion encountering N-terminally oriented JF 646 - labeled CTCF (cyan). Growth of DNA loop (red) stops upon encountering CTCF at timepoints ∼28 – 33 s, and ∼50 – 80 s. DNA loops were visualized by Sytox Orange stain. Scale bar, 2 μm. **d**, Same as (c), but for an event where loop extrusion did not get blocked by CTCF. Growth of the DNA loop continues upon encountering CTCF at the beginning of the kymograph, and CTCF passes into the loop and translocates with it. **e**, Fraction of loop extrusion events blocked upon encountering N- or C-terminally oriented CTCF or dCas9. The force range between 0.04-0.08 pN was best covered in all experiments and was therefore chosen to compare the overall blocking efficiency of N- or C-terminally oriented CTCF or dCas9 (Extended Data Fig. 6c,d). **f**, The DNA tension at the moment of encounter between cohesin and CTCF was calculated by the amount of DNA outside the loop and the DNA end-to-end length (see Methods). **g**, The loop extrusion blocking probability of CTCF when encountered from its N-(blue) and C-terminal side (red) depends on the DNA tension at the moment of encounter. Solid lines are fits of the form 1-exp(-F/F_0_), which are used to compute the force at which 100% blocking is achieved (N-terminal encounters: P_block_(F) = 147(1 -e ^-F / 0.125 pN^); C-terminal encounters: P_block_(F) = 115(1 - e ^-F / 0.357 pN^)). N = 17, 75, 72, 89, 40, 3 per bin for N-terminal encounters. N = 12, 77, 53, 34, 6 per bin for C-terminal encounters.

### CTCF is an orientation-dependent barrier to cohesin diffusing along DNA

Next, we analyzed how CTCF interacts with cohesin that diffuses along DNA. For this purpose, we used an assay in which cohesin associates with DNA in a high-salt resistant manner that is sensitive to cohesin and DNA cleavage ^43^, suggesting that under these conditions cohesin entraps DNA topologically and moves along DNA as has been proposed for cohesive cohesin ^34,44^. We observed that CTCF frequently blocked diffusion of recombinant human cohesin (64±18 %; mean and SD), while the remaining cohesin traversed CTCF multiple times (Fig. 1e-g, Extended Data Fig. 1g-i, Extended Data Fig. 3a-d). In contrast, EcoRI^E111Q^ rarely blocked cohesin (15±3 %; Fig. 1g, Extended Data Fig. 3e). To determine the orientation of the CTCF molecules that had blocked cohesin translocation, we post-labeled the DNA molecules with a marker protein that binds to one of their ends (Extended Data Fig. 3f). This revealed that 75±8 % (mean and SD) of the blocked cohesin complexes faced CTCF’s N terminal side (Extended Data Fig. 1h). This can be attributed to the orientation of CTCF, since inversion of its binding site reversed this blocking behavior (Extended Data Fig. 3g, h). CTCF can thus block cohesin diffusion in a directional manner, suggesting that CTCF contributes to the accumulation of cohesive cohesin at TAD boundaries ^34^.

**Figure 3.**
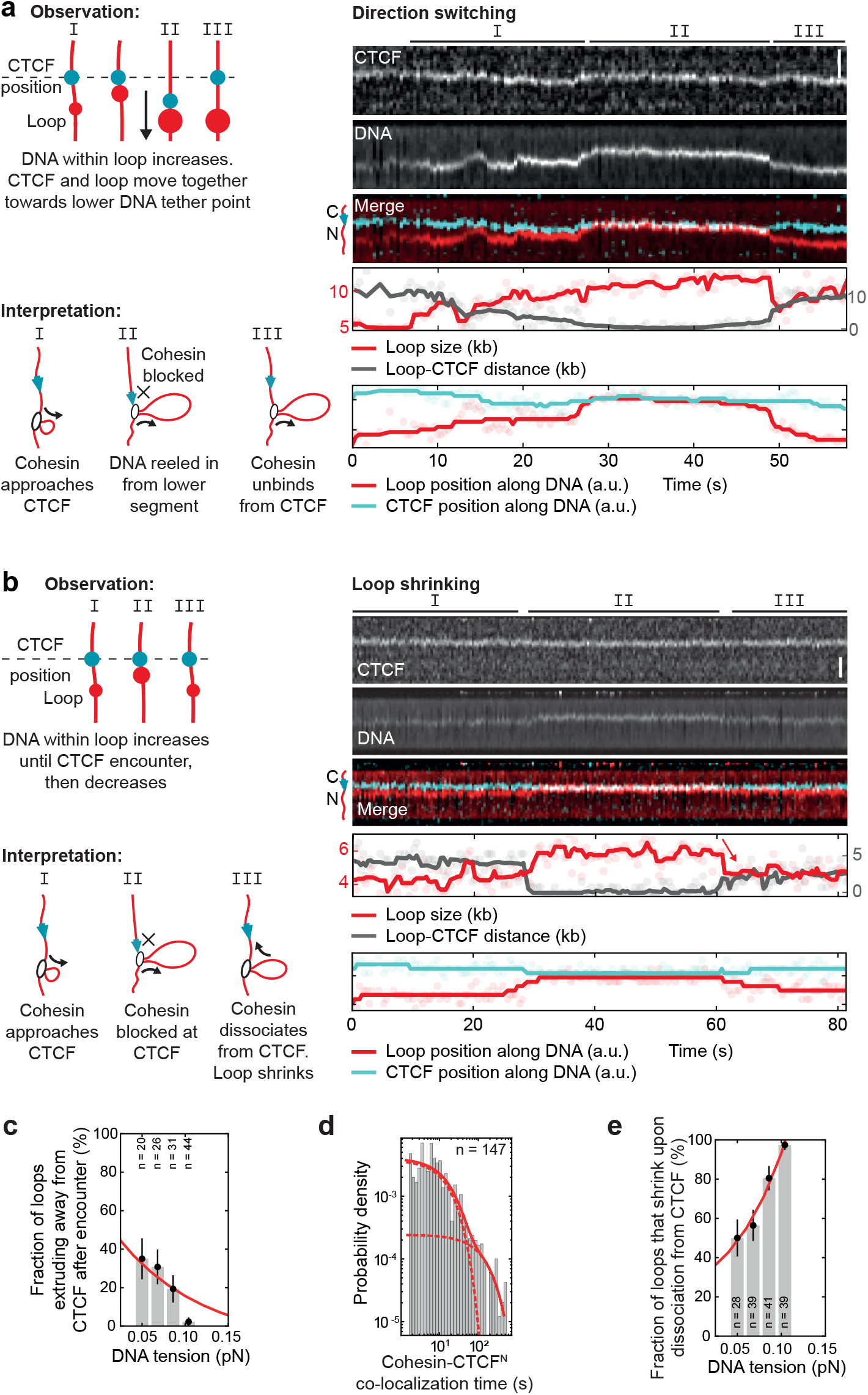
CTCF changes the direction of cohesin-mediated loop extrusion or induces loop shrinkage, depending on the DNA tension. **a, b**, (Left panels) observation and interpretation illustrations of (right panels) kymographs of cohesin-mediated DNA loop extrusion encountering N-terminally oriented, JF646-labeled CTCF (cyan). DNA loops (red) were visualized by Sytox Orange stain. Scale bars, 2 μm. In panel A, the growing loop encounters CTCF at 28 s. CTCF and the growing DNA loop move towards the lower DNA tether point, indicating extrusion on the side facing away from CTCF. In panel B, growth of DNA loop stops upon encountering CTCF at ∼29 s. The DNA l oop shrinks fo l l owing dissociation from CTCF at ∼60 s. **c**, Fraction of loops extruding away from CTCF, versus the DNA tension at the moment of encounter. **d**, Co-localization time of encounters between cohesin and CTCF’s N-terminus. The fit denotes a 2-exponential distribution with rate constants k_1_ = 0.061 s^-1^ and k_2_ = 0.006 s^-1^ (T_1_ ∼ 16 s and T_2_ ∼ 167 s). Dashed l ines represent the individual components of the 2-component exponential distribution; solid line represents the final 2-component exponential distribution. **e**, Fraction of loops which shrink upon release from CTCF versus DNA tension at the moment of encounter.

### CTCF is an orientation-dependent barrier to loop-extruding cohesin

To test whether CTCF also acts as a barrier to loop-extruding cohesin, we introduced a single CTCF site at position 9.7 kb in a 31.8 kb DNA, such that CTCF’s N-terminus would face the longer end of the DNA. We tethered both ends of these molecules to the surfaces of flow cells and stained them with Sytox Orange. We then bound CTCF purified from HeLa cells (Extended Data Fig. 4a – e) to these DNA molecules and introduced HeLa cohesin (Extended Data Fig. 4f), recombinant NIPBL-MAU2 (Extended Data Fig. 1h), and ATP.

**Figure 4.**
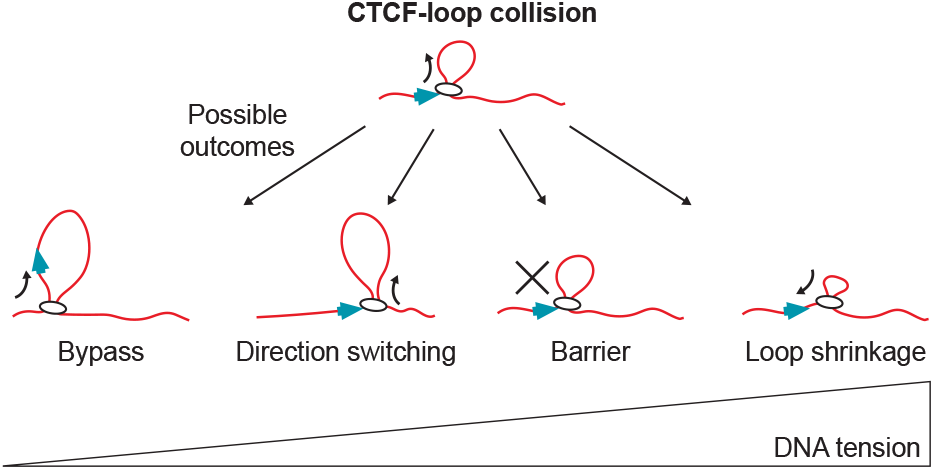
DNA tension affects the outcome of CTCF-cohesin collisions. At low DNA tension, CTCF is frequently incorporated into the growing DNA loop. At higher DNA tensions, CTCF promotes loop-extrusion direction switching, blocks loop extrusion and, at highest DNA tensions, induces loop shrinkage.

Upon buffer flow perpendicular to the DNA axis, CTCF could be detected either at the base of (Fig. 2a) or within DNA loops (Fig. 2b), suggesting that it functioned as a barrier to loop extrusion in some but not all cases. To analyze this behavior quantitatively, we monitored loop extrusion in the absence of buffer flow, where loop formation results in the appearance of a bright spot on the DNA that increases in intensity over time. Tracking and quantification of loop position and size as well as of CTCF position permitted the classification of encounters of cohesin-mediated DNA loops with CTCF (Fig. 2c,d, for additional examples see Extended Data Fig. 5a, b; for Supplementary Videos and animated illustrations, see Supplementary Videos 1 and 2.).

These experiments revealed that N-terminally oriented CTCF blocked the progression of loop extrusion in 45±9 % (mean ± SD) of encounters (Fig. 2c, e, Extended Data Fig. 5a, b), whereas the blocking efficiency was reduced to 16±7 % (mean ± SD) when we used DNA molecules in which the orientation of the CTCF binding site had been inverted and on which cohesin therefore encountered CTCF’s C-terminus (Fig. 2d, e). In contrast, the control protein dCas9, which has a larger mass (180,000) than CTCF-Halo-Flag (118,600) blocked loop extrusion in only 5±10 % (mean ± SD) of encounters (Fig. 2e), consistent with the finding that cohesin can readily traverse non-interacting DNA-bound particles during loop extrusion ^45^.

These results indicate that monomeric CTCF, despite its relatively small mass and Stokes radius (5 nm for CTCF’s N-terminus; ^46^), is sufficient to block loop extrusion by cohesin in a directional manner, possibly because the N-terminus of CTCF can bind cohesin ^26^. Interestingly, the N- and C-terminal blocking frequencies of 45% and 16% observed in our experiments can remarkably well explain *in vivo* estimates of how frequently loops are detected between CTCF sites oriented in a convergent, divergent, or tandem manner (Extended Data Fig. 6a), suggesting that CTCF may be solely responsible for determining how frequently loops are anchored at these differently oriented sites.

### CTCF’s barrier function depends on DNA tension

Unexpectedly, we observed that the CTCF blocking efficiency for loop-extruding cohesin depends on the tension in the DNA that is reeled in. Since DNA molecules are tethered at both ends in our assay, loop extrusion continuously shortens the non-extruded parts of the DNA molecules and thus increases their tension, until this tension exceeds the stalling force of loop extrusion (Fig. 2f and ^11^). We noticed that larger loops and loops extruded from DNAs with a longer end-to-end length tended to be stalled more efficiently by CTCF than those formed from less stretched DNA. Because both scenarios coincide with larger tension in the unextruded part of the DNA, we tethered DNA molecules to the surface of flow cells with various degrees of ‘slack’, performed CTCF loop extrusion blocking assays and calculated the tension that DNA molecules experienced when cohesin encountered CTCF.

Interestingly, the efficiency of CTCF’s barrier activity indeed very strongly correlated with increased DNA tension (Fig. 2g). Strikingly, our data indicate that CTCF does not block loop extrusion by cohesin at all when no force is applied, while CTCF blocks loop extrusion increasingly when tension is applied to the DNA, with CTCF reaching a blocking efficiency of 100 % at ∼0.14 pN. Notably, this tension is close to the ∼0.15 pN force required to stall loop extrusion itself (Extended Data Fig. 6b – d). Encounters from the C-terminal side showed a similar trend, i.e. blocked loop extrusion more frequently at higher tension, but with much lower blocking frequencies. In contrast, the ratio of blocking efficiencies of N-terminal versus C-terminal encounters (3.6 ± 0.8 -fold [mean ± SD]) was unaffected by DNA tension (Extended Data Fig. 6e). These results show that CTCF’s ability to block loop extrusion by cohesin depends on the tension of the DNA that CTCF and cohesin are bound to.

To test whether this phenomenon could relate to cohesin’s possible ability to ‘step over’ CTCF we measured cohesin’s step size during loop extrusion using magnetic tweezers. These measurements showed that cohesin, on average, takes large steps of ∼40 nm (100-200 bp) on DNA and that the step size decreases when DNA tension increases (Extended Data Fig. 7a-d), as also observed for condensin ^47^. We tested in simulations whether cohesin might encounter CTCF more frequently at higher DNA tension because cohesin is less likely to step over CTCF since the extrusion steps become smaller. However, this hypothesis was not supported by our simulations (Extended Data Fig. 7e, f). We therefore suspect that DNA tension increases the blocking efficiency of CTCF by other mechanisms, such as reducing the step frequency at increased tension which allows more time for CTCF-cohesin binding, or decreasing thermal fluctuations of DNA ^48^ which may reduce the space that CTCF has to explore to find cohesin, or that cohesin’s weak motor activity can more easily overcome the low binding affinity of CTCF-cohesin interactions ^26^ at low DNA tension than at high tension (discussed in the legend of Extended Data Fig. 7f). Irrespective of these interpretations, our results indicate that local changes in DNA tension that could be caused by nucleosome assembly, transcription, DNA replication, supercoiling or other processes, can affect genome architecture by modulating the permeability of TAD boundaries.

### CTCF can anchor cohesin loops only for short periods of time

To analyze the fate of loops that were blocked by CTCF, we first analyzed for how long CTCF and loops co-localize under conditions in which the loop size was constant (i.e., where loop extrusion stalled upon encounter). We frequently observed brief (tens of seconds) and repeated encounters between loops and CTCF (Extended Data Fig. 8a, Supplementary Video 3) as well as occasional encounters that lasted for several minutes (Extended Data Fig. 8b). The distribution of CTCF-loop interaction times following stalling events was well-described by a bi-exponential distribution, indicating the existence of two populations with mean CTCF-loop association times of 16 s and 167 s (Fig. 3d, Extended Data Fig. 8c-g).

Unlike CTCF’s blocking function, the CTCF-loop association time was largely unaffected by CTCF orientation (Extended Data Fig. 8c-g). It is conceivable that the infrequent C-terminal blocking events that we observed represent occasions where cohesin in fact encountered CTCF’s N-terminus after passing over its C-terminus. The results indicate that CTCF interacts with cohesin mostly transiently (> 85% of encounters lasted less than 3 minutes; Extended Data Fig. 8g), which is similar to the lifetime that has been measured for particular loops in cells ^16,18^. However, longer-lived loops have been predicted to exist for up to several hours ^32,49^. Since we did not observe such prolonged co-localization of CTCF and loops, additional proteins may be required to anchor loops for such long time periods, for example the PDS5 proteins, which are also required for TAD boundaries in cells ^4^.

### CTCF can switch the direction of cohesin-mediated loop extrusion and shrink loops

It has been speculated that cohesin switches from symmetric to one-sided asymmetric extrusion at TAD boundaries at which loop ‘stripes’ or ‘flames’ have been detected in Hi-C experiments ^23,26,31,32^. We therefore analyzed whether a change in extrusion symmetry could be observed when cohesin encounters CTCF. However, close analysis revealed that, unlike previously assumed ^10,12,13^, cohesin rarely extrudes DNA strictly symmetrically, but instead frequently reels in DNA first from one side and then the other, switching direction multiple times (a detailed analysis of this bidirectional extrusion will be reported in a separate study (Barth et al, in preparation)). We therefore analyzed whether CTCF can trigger a switch of the direction of loop extrusion. To investigate this, we monitored the size of DNA loops and their position relative to CTCF, following encounters that had blocked loop extrusion.

At low DNA tension, we observed events where CTCF indeed switched the direction of cohesin’s loop extrusion activity. Cohesin approached CTCF by reeling in the intervening DNA and then, upon encounter with CTCF, it began to reel in DNA from the other direction while remaining bound to CTCF (Fig. 3a, c, Supplementary Video 4). A control experiment with gold nanoparticles that were tethered to DNA as artificial roadblocks ^45^, reversed the direction of loop extrusion 2.6 times less frequently at low DNA tension (Extended Data Fig. 9a), suggesting that this ability may be a specific property of CTCF. This effect can potentially explain the appearance of ‘stripes’ and ‘flames’ at TAD boundaries.

Interestingly, at higher DNA tension, CTCF did not switch the direction of loop extrusion (Fig. 3c), but instead loops tended to shrink in size upon release from CTCF (Fig. 3b, e, Extended Data Fig. 10, Supplementary Video 5). In most cases, loops decreased in size within a single step (i.e. within the imaging frame speed of 0.4 s, Extended Data Fig. 10a, b, e, f, h), but in some cases loops shrunk gradually over several seconds (Extended Data Fig. 10c-f, h) at a rate similar to that of loop extrusion (Extended Data Fig. 10g). In both cases, loops did not disrupt completely but were reduced in size by several kb and on average lost 35 % of looped DNA (Extended Data Fig. 10i, j). Such loop shrinkage could be observed with similar frequencies when cohesin collided with artificial roadblocks on DNA (Extended Data Fig. 9b), suggesting that this may be a general response of cohesin to encountering barriers on DNA, irrespective of specific binding of the roadblock to cohesin. Its physiological relevance and whether it represents a reversal of the loop extrusion mechanism or ‘slippage’ of DNA from the loop remains to be investigated, but it is interesting to note that the gradual shrinkage occurred at a similar rate as loop extrusion.

## Discussion and conclusions

Our results indicate that CTCF molecules find their cognate binding sites by facilitated diffusion and once bound to them are sufficient as monomers to block passively diffusing cohesin complexes, which possibly reflects how DNA-entrapping cohesive cohesin accumulates at TAD boundaries ^34^. CTCF is also a barrier to actively loop-extruding cohesin, presumably reflecting how CTCF establishes TAD boundaries. As predicted from Hi-C experiments, CTCF performs this function asymmetrically with its N-terminus blocking cohesion almost four-fold more efficiently than at its C-terminus. Unexpectedly, this function is regulated by the tension of the DNA that CTCF and cohesin are bound to, implying that genomic processes that alter DNA tension will modulate the permeability of CTCF boundaries and thus the length of chromatin loops extruded by cohesin.

Our data indicate that encounters with CTCF can alter cohesin’s loop extrusion activity in at least three different ways (Fig. 4): it can block loop extrusion, it can switch its direction, i.e. cause cohesin to reel in DNA from the opposite side as before, and it can lead to a process in which loop formation is reverted as the loop starts shrinking rather than growing. The observation that TADs detected by Hi-C are ‘filled’ with chromatin loops which are not anchored at both TAD boundaries may therefore not only reflect the presence of nascent loops which have not been fully extruded yet, as has been assumed so far, but also the existence of ‘shrunk’ loops which had already reached TAD boundaries but were switched there into a ‘reverse’ mode by CTCF. It is conceivable that such a backtracking process is used as a failsafe mechanism for enabling repeated interactions between specific genomic regions in cases where these remained unproductive upon their first encounter during forward loop extrusion, for example during V(D)J recombination of antigen receptor genes ^5,6^.

Although indirect effects of CTCF on cohesin, for example inhibition of WAPL or promotion of PDS5 binding at the expense of NIPBL, may enhance the establishment of a barrier to loop extrusion as detected in cell-population measurements (reviewed in ^9^), our experiments indicate that these effects are not strictly required. Together, our findings reveal that CTCF controls cohesin and therefore genome architecture via multiple modes. Our results will provide the basis for future mechanistic and physiological studies of CTCF’s key functions in gene regulation, recombination, cell differentiation and tumorigenesis.

## Acknowledgements

We thank D. Goetz for cloning assistance, M. Madalinski for purifying JF646-HaloTag Ligand, T. Lendl for image analysis support, and P. Pasierbek and other members of IMP/IMBA Biooptics facility for assistance.

## Funding

Research in the laboratory of C.D. was supported by ERC Advanced Grant 883684 (DNA looping) NWO grant OCENW.GROOT.2019.012, and the NanoFront and BaSyC programs. Research in the laboratory of J.-M.P. was supported by Boehringer Ingelheim, the Austrian Research Promotion Agency (Headquarter grant FFG-852936), the European Research Council under the European Union’s Horizon 2020 research and innovation programme GA No 693949, the Human Frontier Science Program (grant RGP0057/2018) and the Vienna Science and Technology Fund (grant LS19-029). J.-M.P. is also an adjunct professor at the Medical University of Vienna.

## Author contributions

IFD, RB, MZ, KN, RJ, CD and J-MP designed experiments. IFD performed CTCF loop extrusion experiments, purified proteins, and analyzed data. RB performed dCas9 and stalling force loop extrusion experiments, performed magnetic tweezers experiments, and analyzed data. MZ performed single-molecule CTCF characterization, EMSAs, generated DNA templates for cohesin diffusion experiments, performed cohesin diffusion experiments, purified proteins, and analyzed data. JvdT generated DNA templates for loop extrusion experiments. WT generated the CTCF-Halo-Flag HeLa cell line. KN performed iFRAP experiments and analyzed data. RJ performed magnetic tweezers experiments and analyzed data. JK analyzed magnetic tweezers data. GW purified proteins. IFD and J-MP wrote the manuscript with input from all authors. J-MP and CD supervised the study.

## Competing interests

Authors declare no competing interests.

## Notes

### Competing Interest Statement

The authors have declared no competing interest.

### Summary of Updates

Typographical errors corrected.

