## Supplementary Information for "CTCF is a DNA-tension-dependent barrier to cohesin-mediated DNA loop extrusion"

**Affiliations:**

**This PDF file includes:**

Methods
Methods References
Captions for Supplementary Videos 1 – 5
Extended Data Figures 1 – 10

**Other supplementary material for this manuscript includes:**
Supplementary Videos 1 – 5
Supplementary File 1. DNA construct information.

### Methods

#### DNA constructs for use as substrates in cohesin diffusion assay

DNA fragments containing a single HighOc1 CTCF binding site [50] (TCAGAGTGGCGGCCAGCAGGGGGCGCCCTTGCCAGA) were generated by PCR using Phusion Hot Start DNA polymerase (NEB #M0535S) and inserted into the plasmid pPlat (25,754 bp) at the FspAI (ThermoFisher Scientific #ER1661) restriction site in either forward or reverse complement orientation using Gibson assembly [51]. Constructs were then linearized with the restriction enzyme SpeI (New England Biolabs #R3133S) and biotinylated as previously described [43].

#### DNA constructs for use as substrates in loop extrusion assay

We prepared two constructs of 31.8 kb length which contain a CTCF site placed asymmetrically ~9,7 kb from one end, which allows discrimination of the orientation of the DNA construct based on the binding position of CTCF. One construct was oriented such that the N-terminus of CTCF points to toward the longer end of the DNA (plasmid #121; used for N-terminal encounters) and the motif direction of the other construct was reversed (plasmid #128; used for C-terminal encounters). Plasmid #121 was made of plasmids #64, #66, #67, #69, #118 and #71 (see Supplementary File 1 for a complete list of intermediate vectors and primers used). Plasmid #128 was made of plasmids #64, #66, #124, #69, #118 and #71 (see Supplementary File 1). Plasmids #121 and #128 were cloned using Golden Gate cloning, using BsaI-HFv2 as the type-2 restriction enzyme (NEB E1602). Intermediate vectors (#64, #66, #67, #124, #69, #118 and #71) were made using Gibson assembly and traditional (restriction enzyme based) cloning techniques (see Supplementary File 1) (NEB E2621 Gibson mix; NEB M0515 Q5 polymerase).

Biotin-containing handles were made by a PCR reaction with primers JT337 (biotin-GACCGAGATAGGGTTGAGTG, IDT) and JT338 (biotin-CAGGGTCGGAACAGGAGAGC) on #18 pBluescript SK+ (Stratagene), using GoTaq 2 (Promega, M7845). This results in a 1238 bp PCR fragment, which was cleaned up with Promega Wizard™ SV Gel and PCR Cleanup System (Promega, A9282). Fresh plasmid #121 and #128 were purified using the Qiafilter plasmid midi kit (Qiagen, 12243). After purification, the plasmids were cut with both with XhoI and NotI-HF and biotin handles were cut with either XhoI or NotI-HF. The digested products were mixed together with a ~10x molar excess of the biotin-handle over the linearised plasmid. Ligation was done with T4 DNA ligase (NEB, M0202L), overnight at 16 °C and heat inactivated the next morning for 20 minutes at 65 °C. The resulting 31.8 kb DNA construct was cleaned up using an ÄKTA pure system, with a homemade gel filtration column containing approximately 46 ml of Sephacryl S-1000 SF gel filtration media (Cytiva) in TE + 150mM $NaCl_2$. The sample was run at 0.2 ml/min and fractions of 0.5 ml were collected.

#### DNA constructs for use as substrates in magnetic tweezer assay

DNA constructs for magnetic tweezer experiments of 1.5 kb length were synthesized as described previously [47].

#### DNA constructs for protein expression

Human NIPBL with N-terminal FLAG and Halo tags and a C-terminal 10xHis tag as a tandem construct with untagged human MAU2 in pLib was as described [10]. 6xHis-Halo-EcoRI$^{E111Q}$ and 6xHis-tetR-Halo in pLib were as described [43]. 10xHis-CTCF-Halo-Flag was cloned into pLib by combining the human CTCF ORF and the Halo-tag ORF using Gibson assembly. A C-terminal Flag-tag sequence was introduced as a 5'overhang in the reverse primer used for Halo-tag ORF amplification. To generate 10xHis-CTCF-Halo-Avi-Flag, the 10xHis-CTCF-Halo-Flag vector backbone was amplified around the end of Halo-tag sequence, at which position an Avi-tag was introduced using Gibson assembly.

**Generation of radioactively labelled dsDNA probe for electrophoretic mobility shift assays (EMSA)**

100 bp dsDNA fragments containing wildtype or scrambled versions of the HighOc1 CTCF binding site [50] (wt: TCAGAGTGGCGGCCAGCAGGGGGCGCCCTTGCCAGA) were prepared by overlap extension PCR: two ssDNA oligos with partially overlapping sequences were used in a PCR reaction catalyzed by Phusion Hot Start DNA Polymerase (NEB #M0535S) and purified using a PureLink PCR Purification Kit (Invitrogen #K3110002). 1 pmol of dsDNA probe was subsequently incubated with 0.5 μl [$\gamma$-$^{32}$P]ATP (3000Ci/mmol, 10 mCi/ml; Hartmann Analytic #SCP-301) and T4 Polynucleotide Kinase (PNK) (NEB #M0201S) in a 20 μl reaction at 37 $^{o}$C for 1 hour. T4 PNK was subsequently heat inactivated by incubating the reaction at 65 $^{o}$C for 10 min.

**Generation of methylated dsDNA probe for EMSA**

A 100 bp dsDNA fragment containing the HighOc1 CTCF binding site described above [50] was methylated *in vitro* using M.SssI CpG methyltransferase (NEB #M0226S) following the manufacturer's protocol. To increase methylation efficiency, four rounds of methylation, each followed by DNA purification using PureLink PCR Purification Kit (Invitrogen #K3110002), were performed. Methylation efficiency was assessed by incubating 300 ng of purified methylated DNA with 1 μl of the methylation sensitive restriction enzyme EaeI (NEB #R0508S) in 20 ul reaction containing 1x CutSmart buffer (NEB) at 37oC for 1 hour. The reaction products were resolved by electrophoresis on a 0.8% agarose gel and Ethidum bromide staining was detected using a BioRad ChemiDoc Imaging System. The final dsDNA fragment used as unlabeled, methylated competitor in Fig.1B was methylated with ~80% efficiency.

**Generation of CTCF-Halo-Flag HeLa Kyoto cell line**

HeLa Kyoto cells were cultured as described previously [4]. The CTCF-Halo-Flag cell line was generated by homology-directed repair using CRISPR Cas9 (D10A) paired nickase [52]. A donor plasmid comprising CTCF homology arms (719bp and 459 bp on either side of the coding sequence stop site) and Halo-Flag were cloned into plasmid pJet1.2. Cas9 guide RNA sequences were identified using the website crispr.mit.edu (gRNA1: CACCGCAGCATGATGGACCGGTGA; gRNA2: CACCGGAGGATCATCTCGGGCGTG) and inserted into plasmid pX335 (a gift from Feng Zhang, Addgene, 42335). HeLa Kyoto cells were transfected with donor, Cas9 nickase plasmids using Lipofectamine 2000 (Invitrogen #11668019). 7 days later, cells were labeled with Halo-TMR (Promega #G8251) and sorted by flow cytometry. The clonal cell line was selected after verification of homozygous Halo-Flag insertion by PCR of genomic DNA, immunoblotting and inspection by microscopy.

**Protein expression and purification**

Baculoviruses for protein expression in Sf9 insect cells were generated as described [53]. Expression cultures were incubated at 27 $^{o}$C for 48 – 60 h after infection. Cells were centrifuged, washed in PBS, frozen in liquid nitrogen and stored at -80°C.

Recombinant CTCF protein purification

Baculovirus-infected cell pellets from cultures supplemented with 0.1 mM $ZnCl_2$ were lysed by Dounce homogenization and resuspended in CTCF lysis buffer (35 mM $NaH_2PO_4/Na_2HPO_4$ pH 7.4, 350 mM NaCl, 0.1 mM $ZnCl_2$, 5% glycerol, 0.05% Tween-20, 5 mM Imidazole) supplemented with 1 mM PMSF, EDTA-free cOmplete tablet (1 per 50 ml) (Roche #11873580001), 1 mM DTT and 0.001 U/μl Benzonase. The lysate was cleared by centrifugation at 18,000 g for 1 h at 4$^{o}$C. The soluble fraction was incubated with NiNTA agarose (Qiagen #30230) for 1 h at 4$^{o}$ C and washed with CTCF buffer (35 mM $NaH_2PO_4/Na_2HPO_4$ pH 7.4, 150 mM NaCl, 0.1 mM ZnCl2, 5% glycerol) supplemented with 1 mM DTT and 35 mM Imidazole. For the final washing step, DTT was omitted from the washing buffer. Protein was eluted with CTCF buffer supplemented with 300 mM Imidazole. The eluate was subsequently concentrated approximately two-fold using a Sartorius Vivaspin 50 kDa MWCO concentrator (Sartorius #VS2031) and incubated with M2-FLAG agarose (Sigma Aldrich #A2220) for 90 min at 4˚C. The resin was washed with CTCF buffer and incubated with fluorescent Halo ligand (TMR Promega #G8252; Alexa660 Promega #G8472,) for 15 min at room temperature. Following extensive washing with CTCF buffer, the labelled protein was eluted in CTCF buffer supplemented with 0.5 mg/ml 3xFLAG peptide. The eluate was supplemented with 1 mM DTT, concentrated 2 – 4-fold using a Sartorius Vivaspin 50 kDa MWCO concentrator, flash-frozen and stored at -80$^{o}$ C.

HeLa CTCF-Halo-Flag purification

HeLa CTCF-Halo-Flag protein was purified as described for SCC1-Halo-Flag [10] except 20 mM Tris pH 7.5 was used in all CTCF purification buffers instead of 25 mM $NaH_2PO_4/Na_2HPO_4$ pH 7.5, and 0.1 mM $ZnCl_2$ was included in all purification buffers except for the FLAG elution buffer. HeLa CTCF was labelled with JF 646-HaloTag Ligand.

Recombinant cohesin, HeLa cohesin, NIPBL-MAU2 and EcoRI$^{E111Q}$ protein purification

Recombinant cohesin, HeLa SCC1-Halo-Flag cohesin and recombinant NIPBL-MAU2 were purified as described [10]. EcoRI$^{E111Q}$-Halo and TetR-Halo were purified as described [43].

**Electrophoretic mobility shift assay (EMSA)**

For the competition EMSA assay 60 fmol of recombinant CTCF was mixed with 1 µg poly dI-dC (Thermo Scientific #20148E) in a 20 µl reaction containing 35 mM Tris pH 7.9, 50 mM KCl 50 mM NaCl, 5 mM $MgCl_2$, 0.1 mM $ZnCl_2$, 5% Glycerol, 1 mM DTT and 50 ng/µl BSA at room temperature for 20 min. Subsequently, 21 fmol of [$\gamma$-$^{32}$P]ATP (Hartmann Analytic; SCP-501) labeled dsDNA probe was added in the presence of 100x unlabeled competitors (dI-dC; wild type, scrambled or methylated CTCF oligo), and the reaction was incubated at the room temperature for additional 10 minutes. The binding reactions were loaded on to pre-run (1 hour 100 V, 10 mA, ice-cold water bath, 0.5x TBE running buffer) 4% non-denaturing acrylamide gel and the samples were resolved for 1 hour at the same conditions as the pre-run. The gel was exposed to a Phosphoscreen overnight and analyzed by phosphor imaging with a Typhoon Scanner (GE Healthcare). Images shown are representative of three independent experiments.

**Inverse fluorescence recovery after photobleaching**

CTCF-mEGFP cells [4] were seeded on Lab-Tek chambered coverslips (ThermoFisher Scientific). Before imaging, the culture medium was exchanged for pre-warmed phenol red free medium. To avoid new protein synthesis, Cycloheximide (1 µg/ml) was added before imaging. Imaging was performed with a Zeiss LSM880 confocal microscope using a 40×1.4 Plan-Apochromat DIC oil objective at 37℃ and 5 % $CO_2$. Two images were acquired before bleaching half of the nucleus by 100% transmission of a 488 nm laser. Recovery of fluorescence was recorded over 2 hours at 1 min intervals and normalized to the mean of the pre-bleach fluorescent intensity to the first image after photobleaching and to background. G1 and G2 phase cells were identified by the cellular distribution of DHB-mKate2 signals [54]. Curve fitting was performed with a single exponential function $f(t) = \exp(-k_{off,1}t)$ or double exponential functions $f(t) = a \cdot exp(-k_{off,1}t) + (a-1) \cdot exp(-k_{off,1}t)$ in R using the minpack.lm package (version 1.2.1). Dynamic and stable residence times were calculated from $1/k_{off,1}$and $1/k_{off,2}$, respectively. Double exponential curve fitting was performed under a constraint that $1/k_{off,1}$ and $1/k_{off,2}$ are between 1-30 min and 1-12 hr, respectively. Soluble fractions were estimated by the reduction of fluorescence signals in the unbleached area after photobleaching.

**Recombinant CTCF single molecule imaging characterization**

CTCF flow-in, washing and imaging

Flow cells were incubated with Avidin DN (Vector Laboratories; #A3100) and DNA as described [10] except that pPlat containing single HighOc1 CTCF binding site was used instead of λ-DNA. Flow cells were washed with 400 µl WB buffer (20 mM Tris pH 7.5, 50 mM KCL, 5 mM EDTA) supplemented with 0.1 mg/ml BSA and 10 nM Sytox Green (ThermoFisher Scientific; #S7020) or Sytox Orange (ThermoFisher Scientific; #S11368) at 50 µl/min. 100 µl recombinant CTCF-Halo (labelled with TMR in experiments shown in Fig. 1 and in Extended Data Fig. 1a, b, e, f, i and Extended Data Fig. 3; or labelled with Alexa 660 in experiments shown in Extended Data Fig. 1c, d) was then introduced into the flow chamber at 2.5 nM final concentration in CL100 buffer (35 mM Tris pH 7.5, 100 mM KCL, 5 mM $MgCl_2$, 5% glycerol, 0.005% Tween-20, 0.1 mg/ml BSA, 1 mM TCEP) at 30 µl/min and subsequently incubated for 4 min without buffer flow. Flow cells were then washed with CL150 buffer (CL100 buffer supplemented with 50 mM KCl) at rate of 50 µl/min to remove non-specifically bound CTCF molecules.

To determine the orientation of DNA molecules following image acquisition, TMR labelled EcoRI$^{E111Q}$-Halo or TetR-Halo was flowed into flow cells at 2 nM or 5 nM final concentration, respectively, in EcoRI buffer (20 mM Tris 7.5, 150 mM KCl, 0.1 mg/ml BSA) supplemented with 10 nM Sytox Green at 30 µl/min, incubated for 4 min and washed with 200 ul of EcoRI buffer.

All recombinant CTCF single molecule imaging characterization and cohesin diffusion assay experiments were performed at room temperature. Unless stated otherwise, time-lapse microscopy images were acquired at 4 s intervals using a Zeiss TIRF 3 Axio Observer setup and 488 nm, 561 nm and 639 nm lasers [43]. A protocatechuic acid (PCA)/protocatechuate-3,4-dioxygenase (PCD)/Trolox oxygen scavenger system (final concentration 10 nM PCD, 2.5 mM PCA, 2 mM Trolox); was added to all buffers used during data acquisition.

Imaging the kinetics of recombinant CTCF association with DNA

To image the kinetics of CTCF association with DNA (Fig. 1c, d, Extended Data Fig. 1a), 0.5 nM TMR labelled CTCF-Halo was introduced into flow cells in CL100 buffer at 30 µl/min. For experiments shown in Fig. 1d and Extended Data Fig. 1a, images were acquired at 3.12 sec/frame intervals. For measurements of CTCF residence time on DNA (Fig. 1c, Extended Data Fig. 1e-f) images were acquired at 10.15 sec/frame intervals.

Positional analysis of recombinant CTCF on DNA

The position of recombinant CTCF on DNA was analyzed in Fiji. EcoRI or TetR mediated end-labeling was used to unambiguously assign the orientation of DNA strands tethered to the surface. The distance between the center of the mass of fluorescent intensity signal marking the DNA end and the fluorescent signal of protein was measured, and the ratio between measured distance and the total length of DNA molecule was calculated as a position along the DNA in bp. Single-molecule tracking of CTCF position was performed using custom Fiji macro.

Recombinant CTCF photobleaching analysis

To quantify the number of recombinant Alexa 660 (A660) labeled CTCF molecules bound at a CTCF DNA binding site, A660 signals on DNA were identified in laser profile corrected images, subtracted from the local background, averaged over 10 frames and plotted in Extended Data Fig. 1d.

**HeLa CTCF single molecule imaging characterization**

CTCF flow-in, washing and imaging

Flow cells [43] were incubated with 1 mg/ml Avidin DN (Vector Laboratories) for 15 min and washed extensively with DNA buffer (20 mM Tris pH 7.5, 150 mM NaCl, 0.25

mg/ml BSA (ThermoFisher Scientific; AM2616)). 150 µl of 31.8 kb DNA containing a single CTCF site and biotinylated ends was introduced into flow cells at ~20 pM final concentration at 50 µl/min in DNA buffer supplemented with 20 nM Sytox Orange (ThermoFisher Scientific; S11368). Flow cells were washed with 400 µl of wash buffer 2 (50 mM Tris pH 7.5, 50 mM NaCl, 2.5 mM $MgCl_2$, 0.25 mg/ml BSA, 0.05% Tween-20, 20 nM Sytox Orange) at 100 µl/min, followed by 100 µl of imaging buffer (50 mM Tris pH 7.5, 50 mM NaCl, 2.5 mM $MgCl_2$, 0.25 mg/ml BSA, 0.05% Tween-20, 0.2 mg/ml glucose oxidase (Sigma; G2133), 35 mg/ml catalase (Sigma; C- 40), 9 mg/ml b-D-glucose, 2 mM Trolox (Cayman Chemical; 10011659)) and 5 mM ATP (Jena Biosciences; NU- 1010-SOL)) supplemented with 20 nM Sytox Orange at 100 µl/min. Stock solutions of glucose oxidase (20 mg/ml), catalase (3.5 mg/ml) and glucose (450 mg/ml) were prepared as described [55]. JF646 labelled HeLa CTCF was then introduced into the flow chamber at 0.5 nM final concentration in 100 µl imaging buffer supplemented with 20 nM Sytox Orange at 30 µl/min. Non-specifically bound CTCF was removed by washing three times with 100 µl imaging buffer supplemented with 220 nM Sytox Orange at 100 µl/min.

All HeLa CTCF single molecule characterization and loop extrusion experiments were performed at 37 °C. Time-lapse microscopy images were acquired using a Zeiss Elyra 7 with Lattice SIM$^2$ equipped with 561 nm and 639 nm lasers, two PCO Edge 4.2 sCMOS cameras and a 63×/1.46 Alpha Plan-Apochromat oil objective. 100 ms exposure time images were acquired sequentially for each channel at 0.4 s intervals in HILO mode.

<u>HeLa CTCF photobleaching analysis</u>

To quantify the number of HeLa JF646 labelled CTCF molecules bound at a CTCF DNA binding site, JF646 signals on DNA were identified in laser profile corrected images acquired, subtracted from the local background and averaged over all frames prior to a bleaching event and plotted in Extended Data Fig. 4d. The number of bleaching steps per molecule was determined manually and indicated on Fig. S4E. The fluorescence intensity of molecules bound at a CTCF DNA binding site that bleached in a single step was 2.2 ± 0.6 (mean ± SD).

<u>HeLa CTCF positional analysis</u>

The position of HeLa CTCF was analyzed as described in the section 'Determination of DNA loop size and position of single molecules' below.

**Cohesin diffusion assay and image analysis**

Cohesin diffusion assays were performed essentially as described [43]. CTCF was introduced into flow cells at 2 nM final concentration and incubated for 4 minutes as described in the Recombinant CTCF imaging section above. Flow cells were then washed with CL150* buffer (35 mM Tris pH 7.5, 75 mM NaCl, 75 mM KCl, 1 mM $MgCl_2$, 10% glycerol, 0.003% Tween-20, and 0.1 mg/ml BSA). Cohesin and NIPBL-MAU2 were introduced into flow cells at 0.8 – 2 nM and 2 nM, respectively in 100 µl of CL100* buffer (35 mM Tris, pH 7.5, 50 mM NaCl, 50 mM KCl, 1 mM $MgCl_2$, 10% glycerol, 0.003% Tween-20, and 0.1 mg/ml BSA) at 30 µl/min. Flow cells were incubated for a further 4 min without buffer flow and then washed with CL250* buffer (35 mM Tris pH 7.5, 125 mM NaCl, 125 mM KCl, 1 mM $MgCl_2$, 10% glycerol, 0.003% Tween-20, and 0.1 mg/ml BSA). Cohesin and CTCF imaging was then performed in the absence of buffer flow for 160 sec at 4 sec/frame intervals. Image acquisition was repeated for 3 – 5 fields of view. DNA orientation was determined by flowing in Sytox Green and EcoRIE111Q-Halo or TetR-Halo as described in the Recombinant CTCF imaging section above. Biotin-conjugated quantum dots QD705 (Invitrogen, #Q101163MP) or CTCF-Halo-Avi-Biotin were used as fiducial markers.

CTCF-cohesin channels were aligned with TetR/EcoRI$^{E111Q}$-DNA channels using a custom-written Fiji macro. Each DNA molecule containing diffusing cohesin was manually

examined for the presence of a single CTCF signal positioned at the regions were the CTCF binding site was introduced. DNA molecules containing multiple or non-specifically bound CTCF molecules were excluded from the analysis. The number of diffusing cohesin foci on the selected DNA molecules was determined and DNA molecules containing more than 4 mobile cohesin foci were excluded from the analysis. Cohesin behaviour on DNA was then analysed and classified as follows:
1) Cohesin diffusion blocked: a) cohesin diffuses freely along the DNA and reaches CTCF roadblock, bounces back but does not go past the roadblock during the time of imaging. b) cohesin diffuses freely along the DNA, reaches CTCF and becomes immobilized. c) two or more cohesin molecules blocked by CTCF.
2) Cohesin passes CTCF in one direction: cohesin passes CTCF during imaging and diffuses back towards CTCF but does not pass back to the other side.
3) Cohesin passes CTCF multiple times.
DNAs with the following events were also excluded from analysis: a) Cohesin diffusing or co-localizing with CTCF. b) Cohesin failing to encounter CTCF. c) Cohesin blocked by a high fluorescence intensity CTCF signal, likely a multimer. d) Cohesin or CTCF bleaches during image acquisition.

**Loop extrusion assay**

Perpendicular flow loop extrusion assays were performed essentially as described [10,55]. Flow cells were incubated with 1 mg/ml Avidin DN (Vector Laboratories) for 15 min and washed extensively with DNA buffer (20 mM Tris pH 7.5, 150 mM NaCl, 0.25 mg/ml BSA (ThermoFisher Scientific; AM2616)). 40 µl of 31.8 kb DNA containing a single CTCF site and biotinylated ends was introduced into flow cells at ~ 3 pM final concentration at 15 µl/min in DNA buffer supplemented with 20 nM Sytox Orange (ThermoFisher Scientific; S11368). Flow cells were washed with 20 µl of wash buffer 1 (50 mM Tris pH 7.5, 200 mM NaCl, 1 mM $MgCl_2$, 5% glycerol, 1 mM DTT, 0.25 mg/ml BSA, 20 nM Sytox Orange) at 5 µl/min. Flow was then switched to perpendicular mode and a further 350 µl of wash buffer 1 was introduced at 100 µl/min. 400 µl of wash buffer 2 (50 mM Tris pH 7.5, 50 mM NaCl, 2.5 mM $MgCl_2$, 0.25 mg/ml BSA, 0.05% Tween-20, 20 nM Sytox Orange) was then introduced at 100 µl/min, followed by 100 µl of imaging buffer (50 mM Tris pH 7.5, 50 mM NaCl, 2.5 mM $MgCl_2$, 0.25 mg/ml BSA, 0.05% Tween-20, 0.2 mg/ml glucose oxidase (Sigma; G2133), 35 mg/ml catalase (Sigma; C- 40), 9 mg/ml b-D-glucose, 2 mM Trolox (Cayman Chemical; 10011659)) and 5 mM ATP (Jena Biosciences; NU- 1010-SOL)) supplemented with 20 nM Sytox Orange at 100 µl/min. JF646 labelled CTCF was then introduced into the flow chamber at 0.5 nM final concentration in 100 µl imaging buffer supplemented with 20 nM Sytox Orange at 30 µl/min. Non-specifically bound CTCF was removed by washing three times with 100 µl imaging buffer supplemented with 220 nM Sytox Orange at 100 µl/min. HeLa cohesin and recombinant NIPBL-MAU2 were then introduced into the flow chamber at 0.5 nM and 3.54 nM, respectively, in 250 µl imaging buffer supplemented with 220 nM Sytox Orange at 30 µl/min.

For loop extrusion assays in the absence of buffer flow, flow cells were incubated with Avidin DN and washed with DNA buffer as above. DNA was introduced at 15 µl/min – 25 µl/min to vary the DNA tension. Flow cells were then washed and incubated as above without switching to perpendicular mode.

**dCas9 binding to DNA**

crRNA sequences were chosen at roughly 1/3 of the DNA length and at each end, two sequences were used for efficient binding of the dCas9-gRNA complex per DNA. If located at the same ends, crRNA sequences were spaced at least 2 kb apart to allow discrimination (additionally to bleaching curves) of occasional binding of two dCas9-gRNA complexes per

DNA end. Binding sequences were chosen using CRISPOR (http://crispor.tefor.net/crispor.py; PAM indicated in bold):

seq7932: ACTGGACTGCGACCGGGCAG**GGG**
seq11802: CGCGGTGGAGGCAGACGTGG**CGG**
seq18967: CTGGTTATGCAGGTCGTAGT**GGG**
seq21005: GGCATACAAATATTCCATGA**AGG**

gRNA was obtained by annealing a mixture of Alt-R Crispr-Cas9 ATTO550-labeled tracrRNA and crRNA (IDT) at 95 °C for 2.5 min and slow cooling to 5 °C in steps of 5 °C for 2.5 min each. To couple gRNA to dCas9, 200 nM dCas9 (NEB; NEBM0652T) was mixed with 2 µM gRNA on ice in NEBuffer3.1, incubated at 37 °C for 10 min and placed on ice again.

To bind the dCas9-gRNA complex to DNA, DNA constructs of 31.8 kb length were used to facilitate measurements at a similar end-to-end length and force regime as for CTCF experiments. DNA was bound to the pegylated glass surface and unbound DNA was washed off with 100 µl imaging buffer. Then, 1 nM dCas9-gRNA was flushed in the flow cell and incubated for 5 min. Non-specifically bound dCas9-gRNA was removed by flushing with 100 µl imaging buffer, supplemented with 1 mg/ml heparin. Heparin was removed by washing with 100 µl imaging buffer. This typically left one to two dCas9-gRNA complexes per DNA. Loop extrusion experiments were then performed as described above with 30 pM cohesin and 75 pM NIPBL-Mau2. DNA was visualized by staining with 25 nM SytoxGreen and exciting by a 488 nm laser. gRNA-ATTO550 was excited by 561 nm laser light in an alternating excitation scheme using 60x oil immersion, 1.49NA CFI APO TIRF (Nikon) objective. Emission was collected on a Photometrics Prime BSI sCMOS camera using continuous imaging and an exposure time of 100 ms per frame.

**Loop extrusion image analysis**

Fluorescence images were cropped to regions of interest spanning single stretched DNA molecules. Cropped images were then further analysed in a custom written python-based as described [45] which allows semi-automated processing and inspection of the data. Images were median-filtered and kymographs were constructed along the long axis of the stretched DNA molecule.

<u>End-to-end length of DNA</u>

The ends of the DNA molecule were determined by a 'peak peeling' algorithm: a time span of the kymograph in which no loop extrusion occurs (usually before flushing in of cohesin) is selected and temporally averaged, which yields an intensity profile of DNA along its long axis. Gaussian peaks of width roughly equal to the full-width-half-maximum (FWHM) of the microscope's point spread function (here 300 nm) were placed at the position of maximum intensity. The height of the Gaussian equals the maximum intensity of the profile. The Gaussian was then subtracted from the intensity profile. The same procedure was iteratively applied on the new profile until less than 10 % of the original area under the intensity profile remained. The location of the outermost peaks corresponded to the ends of the DNA and the difference between the peaks corresponded to the end-to-end length.

<u>Determination of DNA loop size and position of single molecules</u>

Peaks in each frame corresponding to DNA loops or single molecules (CTCF or dCas9, from here on referred to as CTCF) were detected in every frame, if present, using the *scipy.find_peaks* algorithm [56]. Detected spots were then connected using the *trackpy* package [57] which allows for tracking of loop and single molecule positions over time. The intensity of the loop ($I_{loop}$) is computed as the summation of 7 pixels surrounding the peak position, corrected for the amount of DNA outside of the loop that falls within this window. The size of the loop $L_{loop}$ is computed by the fraction of intensity attributed to the loop, in relation to the total intensity along the entire DNA molecule and the DNA length $L = 31.8\ kb$,

$$L_{loop} = \frac{I_{loop}}{I} \cdot L.$$

The position of the loop $x_{loop}$ (in bp) is reported relative to one randomly chosen DNA end and thus depends on the DNA intensity between the chosen end to the CTCF (the lead DNA; $I_{lead}$),

$$x_{loop} = \frac{I_{lead}}{I} \cdot L.$$

The loop-CTCF distance is computed similarly by measuring the DNA intensity between loop and CTCF, $I_{loop-CTCF}$. The integration of the DNA intensity between loop and CTCF position was corrected for half of the loop size since loops appear as round foci and overlap the DNA connecting loop and CTCF. The loop-CTCF distance is then

$$d_{loop-CTCF} = \frac{I_{loop-CTCF}}{I} \cdot L.$$

The binding position of CTCF was measured relative to the DNA's end points as determined by peak peeling (see above). Given the lower ($x_{low}$) and upper ($x_{up}$) DNA end points, the tracked CTCF position in space, $x_{CTCF}$, is converted to its sequence position $L_{CTCF}$ as

$$L_{CTCF} = \frac{x_{CTCF} - x_{low}}{x_{up} - x_{low}} \cdot L.$$

MSD calculation

From the position of the CTCF over time, the MSD is computed over a moving window, whose length $w$ was roughly adjusted based on the time before and after encounter of the CTCF by cohesin (usually between 21 and 51 frames, corresponding to ~8 s to ~20 s) as

$$\mathrm{MSD}(t) = \frac{1}{w} \sum_{\tau=t-\frac{w}{2}}^{t+\frac{w}{2}} |x_{CTCF}(\tau + \Delta t) - x_{CTCF}(\tau)|^2,$$

Force calculation

The DNA tension within the loop is zero in the absence of buffer flow. In the fraction of DNA outside the loop, the tension strongly depends on the end-to-end length of the DNA. The amount of DNA outside the loop, $L_{non-loop}$ is computed by subtracting the loop size from the length of the DNA construct. Its contour length is computed as

$$C_{non-loop} = \alpha I_{non-loop} \cdot 0.342 \text{ nm},$$

accounting for the distance between base pairs of 0.324 nm and $\alpha$ is a correction factor thereof to account for the slightly different contour length of DNA molecules when bound by Sytox Orange [58]. Values of $\alpha$ were measured by generating force-extension curves of 10 kb DNA constructs using Magnetic Tweezers [47] and fit to the worm-like chain (WLC) model which was already later used to compute the DNA tension [59] (see below; Supplementary Table 1).

**Supplementary Table 1**: Correction factor $\alpha$ and persistence length $L_p$ for varying concentrations of Sytox Orange (SxO) and Sytox Green (SxG).

| SxO/SxG concentration [nM] | Correction factor $\alpha$ | Persistence length $L_p$ [nm] |
|---|---|---|
| 0 | 1 | 46.1 |
| 10 | 1.0258 | 41.9 |
| 50 | 1.0523 | 36 |
| 100 | 1.0649 | 35.1 |
| 200 | 1.0948 | 37.1 |
| 500 | 1.3829 | 37.2 |

The relative extension of the DNA outside of the loop is computed as

$$f = \frac{C_{non-loop}}{x_{up} - x_{down}}.$$

Together with the persistence length of DNA at the respective concentration of Sytox Orange (Supplementary Table 1), the tension acting on the DNA molecule is then computed according to [59]:

$$F = \frac{k_B T}{L_p}\left(\frac{1}{4 \cdot (1 - f)^2} - \frac{1}{4} + \sum_{i=1}^{7} a_i f^i\right),$$

where $k_B T = 1.3806503 \cdot 10^{-23} \cdot 295\ K \cdot 10^{-18}\ pN \cdot \mu m$ and the coefficients are $a_1 = 1, a_2 = -0.5164228, a_3 = -2.737418, a_4 = 16.07497, a_5 = -38.87607, a_6 = 39.49944, a_7 = -14.17718$.

Smoothing of loop size, loop-CTCF distance and MSD traces

Traces of loop size and loop-CTCF distance were filtered by an edge-preserving Chung-Kennedy filter [60] with $N = 4, 8, 16$, and 32 sampling window lengths, weighting parameter $M = 10$ and sharpness parameter $P = 40$ or a median filter with window size 21 frames.

CTCF-loop co-localization analysis

The duration of CTCF-loop co-localization was identified on smoothed (Chung-Kennedy-filtered) time traces of loop-CTCF distances. Connected stretches at which the loop and CTCF colocalize ($d_{loop-CTCF} = 0$) were identified and components with a gap of only one frame were merged to avoid over-segmenting connected components due to noise in images and erroneous tracking of loop and CTCF positions.

Determination of loop shrinkage rate, time span and size of slipped loop

Dissociation events were identified by the time point at which the loop-CTCF distance increased above 0 pixels. To avoid confounding effects of loop release with subsequent re-initiation of loop extrusion, the end point of the loop shrinkage period was marked manually at the time point at which the loop stops shrinking. The loop size before the slippage event was computed as an average over the loop size in the 5 frames (2 seconds) before the slippage event (over which the loop size remained constant). The loop size after the slipping event was based on the average of 1-5 frames, depending on if the loop immediately grew again after slipping (then only the last frame in which the loop still decreases in size can be used) or retained its size (then the loop size was averaged over up to 5 frames). The difference between the loop size before and after slipping corresponds to the fraction of the loop which was lost upon dissociation. The loop shrinkage rate was determined by a linear fit of the loop size between the dissociation time point between loop and cohesin and the time point at which the looped size stopped decreasing.

Fitting of CTCF-loop co-localization time distributions

Distributions of CTCF-loop co-localization times for N- and C-terminal encounters were fit to mono-, bi-exponential and lognormal distributions by a Maximum Likelihood Estimation routine using *scipy.optimize.minimize* [56]. The log-likelihood log $(L)$ and the number of model parameters $k$, as well as the number of data points $n$ was used to compute the Bayesian information criterion (BIC) for each of the models and distributions as displayed in Figure S8C: BIC $= k \cdot log(n) - 2L$.

Measuring the stalling force of cohesin

The stalling force of cohesin was measured on DNA molecules without binding protein (i.e. without CTCF, dCas9 or EcoRI) which were stretched to > 50% of its contour length (contour length ~11.6 µm for a DNA length of 31.8 kb in imaging buffer containing 100 nM Sytox Orange). On such stretched molecules, the stalling force is reached before the cohesin reaches one end of the DNA, thus eliminating confounding effects from the DNA ends. For kymographs with multiple extrusion, slipping and direction change events (as e.g. in Extended Data Fig. 6a,b), the highest force value (as long as it did not approach one of the DNA ends) during the acquisition time was considered as the stalling force. Note that, in contrast to the measured stalling forces on the fluorescence single-molecule assay, cohesin-mediated loop extrusion steps, measured using Magnetic Tweezers (see below) are still present at considerably higher forces, up to 1 pN. However, only single or a few steps can be observed. So few steps likely accumulate to loop sizes ≤ 1 kb and are thus not visible in the fluorescence assay. Additionally, cohesin molecules may stochastically halt loop extrusion even on DNA with low end-to-end distance and thus at very low tension (see e.g. Extended Data Fig. 6a, timepoint 60 s). Thus, a mixture of cohesin molecules is measured, some of which reach their stalling force and therefore stop extruding, and some of which halt loop extrusion before for other reasons. Therefore, the stalling force shown in Extended Data Fig. 7b is thus a conservative estimate of the stalling force, but comparable to encounters with CTCF and dCas9 since these data points were acquired under identical conditions.

**Magnetic Tweezers experiments**

The magnetic tweezers instrument and experiments were conducted essentially as described [47] with minor modifications. The instrument consisted of a pair of vertically aligned, 1 mm apart, permanent neodymium-iron-boron magnets (Webcraft GmbH, Germany) that were was used to generate the magnetic field [61]. The magnet pair was placed on a motorized stage (translation: Physik Instrumente #M-126.PD2; rotation: Physik Instrumente #C-150.PD) and the light of a red LED ($\lambda$ = 630 nm) was allowed to pass the magnet pair gap in order to illuminate the sample. Transmission was collected by a 50x oil-immersion objective (CFI Plan 50XH, Achromat; NA = 0.9, Nikon), and the bead diffraction patterns were recorded with a 4-megapixel CMOS camera (Falcon 4M60; Teledyne Dalsa, USA) at 50 Hz. The real-time tracking of the magnetic bead movement in all three dimensions was conducted with a LabView 2011-based (National Instruments, USA) control software described and published in [62,63]. Surface-adhered 1.5 µm polystyrene reference beads (PolySciences, Germany) were used as reference to correct for instrumental drift occurring during measurements. 100-200 beads could be tracked simultaneously in one field of view with a spatial resolution of ~2 nm for the used 1.5 kbp long dsDNA tethers [47].

The flow cell and DNA tethering were prepared as described previously [47]. In brief, the reference beads were diluted 1:1500 in PBS buffer (pH 7.4; Sigma Aldrich, USA) and then adhered (~5 min) to the cover glass surface of the flow cell. After washing non-adhered beads out with PBS, sheep digoxigenin antibodies (Roche, Switzerland) at a concentration of 0.1 mg/ml was incubated for 1 h within the flow cell, following a 500 µl washing with PBS and 2 h incubation of 10 mg/ml BSA (New England Biolabs, UK) diluted in PBS (pH 7.4) buffer. After washing with 500 µl PBS buffer, 1 pM of the 1.5 kbp linear dsDNA construct was incubated in PBS buffer for 20 min in the flow cell. After washing again with 500 µl

PBS buffer, streptavidin-coated superparamagnetic beads (DynaBeads MyOne, LifeTechnologies, USA; prior diluted 1:400 from stock in PBS buffer) with a diameter of 1 µm were added resulting in the attachment of the beads to the surface-tethered dsDNA constructs after ~5 min; unbound beads were washed out afterwards with PBS buffer.

Prior to cohesin loop extrusion experiments, the quality of tethered dsDNA constructs was assessed by applying a combination of zero and high force (8 pN), and 30 rotations to each direction at high force: only tethers with singly bound dsDNA and correct DNA end-to-end lengths were used for the subsequent single-molecule experiments. After washing the flow cell with cohesin reaction buffer (40 mM Tris-HCl pH 7.5, 50 mM NaCl, 2.5 mM $MgCl_2$, 1 mM DTT, 0.25 mg/ml BSA, 0.05% Tween-20), 0.1 nM cohesin and 0.25 nM NIPBL-Mau2 were introduced in cohesin buffer supplemented with 2 mM ATP to stretched dsDNA tethers at high force (8 pN). For force-titration experiments (Extended Data Figure 7), the force was lowered in individual experiments to 1, 0.8, 0.6, 0.4, 0.3, 0.2, and 0.1 pN, and maintained for 10 min. All Magnetic Tweezer experiments were performed at room temperature.

The *Z*-bead position over time was extracted with custom-written scripts in IGOR Pro (v6.37, Wavemetrics, USA), as previously described [47,64] and a custom-written automated step detection algorithm (MatLab, MathWorks, USA) was applied to the individual traces as described [47,65] in order to extract individual loop extrusion step sizes. Step sizes measured at the same condition from different traces and experiments were pooled and converted into basepairs [47] to construct the distribution of cohesin step sizes in dependence of force (Extended Data Fig. 7c,d).

**Simulating the encounter probability of cohesin and CTCF, given force-dependent cohesin step sizes**

A 10 kb stretch of DNA was simulated on which CTCF was assumed to be positioned 7 kb from one end. The cohesin binding site was uniformly sampled along the DNA length. For each force value, step sizes were sampled from the empirically obtained distribution as measured by Magnetic Tweezer experiments. The simulations were repeated 500 times for every force value and events in which cohesin came within 50 bp of CTCF were counted as encounters, which constitutes a conservative threshold for the interaction distance between cohesin and CTCF.

**Computation of the combinatorial probability of CTCF-CTCF loops**

The probability to observe CTCF-CTCF loops with the two respective CTCF sites in a convergent (><), tandem (>> and <<), or divergent (<>) manner, as reported previously from Hi-C data [4,66-68], was obtained from the loop extrusion stalling probability upon encounter with CTCF on its N-terminal ($P^N$) and C-terminal ($P^C$) side in the force range 0.04-0.08 pN, as shown in Figure 2e:

$$P(><) = P^N P^N / A$$
$$P(\gg \vee \ll) = P^N P^C / A$$
$$P(<>) = P^C P^C / A,$$

where $A$ is a normalization constant, i.e.

$$A = P(><) + P(\gg \vee \ll) + P(<>).$$

**Data availability**

Original imaging data, DNA constructs and cell lines are available upon request.

**Code availability**

All code used in this study is available upon request.

### Methods References


50 Plasschaert, R. N. *et al.* CTCF binding site sequence differences are associated with unique regulatory and functional trends during embryonic stem cell differentiation. *Nucleic Acids Res* **42**, 774-789, doi:10.1093/nar/gkt910 (2014).

51 Gibson, D. G. *et al.* Enzymatic assembly of DNA molecules up to several hundred kilobases. *Nat Methods* **6**, 343-345, doi:10.1038/nmeth.1318 (2009).

52 Ran, F. A. *et al.* Double nicking by RNA-guided CRISPR Cas9 for enhanced genome editing specificity. *Cell* **154**, 1380-1389, doi:10.1016/j.cell.2013.08.021 (2013).

53 Weissmann, F. & Peters, J. M. Expressing Multi-subunit Complexes Using biGBac. *Methods Mol Biol* **1764**, 329-343, doi:10.1007/978-1-4939-7759-8_21 (2018).

54 Spencer, S. L. *et al.* The proliferation-quiescence decision is controlled by a bifurcation in CDK2 activity at mitotic exit. *Cell* **155**, 369-383, doi:10.1016/j.cell.2013.08.062 (2013).

55 Bauer, B. W. *et al.* Cohesin mediates DNA loop extrusion by a "swing and clamp" mechanism. *Cell* **184**, 5448-5464.e5422, doi:10.1016/j.cell.2021.09.016 (2021).

56 Virtanen, P. *et al.* SciPy 1.0: fundamental algorithms for scientific computing in Python. *Nat Methods* **17**, 261-272, doi:10.1038/s41592-019-0686-2 (2020).

57 Allan, D., Caswell, T., Keim, N., van der Wel, C. M. & Verweij, R. soft-matter/trackpy: Trackpy v0. 5.0. *Genève: Zenodo* (2021).

58 Biebricher, A. S. *et al.* The impact of DNA intercalators on DNA and DNA-processing enzymes elucidated through force-dependent binding kinetics. *Nat Commun* **6**, 7304, doi:10.1038/ncomms8304 (2015).

59 Bouchiat, C. *et al.* Estimating the Persistence Length of a Worm-Like Chain Molecule from Force-Extension Measurements. *Biophysical Journal* **76**, 409-413, doi:https://doi.org/10.1016/S0006-3495(99)77207-3 (1999).

60 Chung, S. H. & Kennedy, R. A. Forward-backward non-linear filtering technique for extracting small biological signals from noise. *J Neurosci Methods* **40**, 71-86, doi:10.1016/0165-0270(91)90118-j (1991).

61 Lipfert, J., Hao, X. & Dekker, N. H. Quantitative modeling and optimization of magnetic tweezers. *Biophys J* **96**, 5040-5049, doi:10.1016/j.bpj.2009.03.055 (2009).

62 Cnossen, J. P., Dulin, D. & Dekker, N. H. An optimized software framework for real-time, high-throughput tracking of spherical beads. *Rev Sci Instrum* **85**, 103712, doi:10.1063/1.4898178 (2014).

63 De Vlaminck, I. *et al.* Mechanism of homology recognition in DNA recombination from dual-molecule experiments. *Mol Cell* **46**, 616-624, doi:10.1016/j.molcel.2012.03.029 (2012).

64 Janissen, R. *et al.* Global DNA Compaction in Stationary-Phase Bacteria Does Not Affect Transcription. *Cell* **174**, 1188-1199.e1114, doi:10.1016/j.cell.2018.06.049 (2018).

65 Loeff, L., Kerssemakers, J. W. J., Joo, C. & Dekker, C. AutoStepfinder: A fast and automated step detection method for single-molecule analysis. *Patterns (N Y)* **2**, 100256, doi:10.1016/j.patter.2021.100256 (2021).

66 de Wit, E. *et al.* CTCF Binding Polarity Determines Chromatin Looping. *Mol Cell* **60**, 676-684, doi:10.1016/j.molcel.2015.09.023 (2015).

67 Guo, Y. *et al.* CRISPR Inversion of CTCF Sites Alters Genome Topology and Enhancer/Promoter Function. *Cell* **162**, 900-910, doi:10.1016/j.cell.2015.07.038 (2015).

68 Haarhuis, J. H. I. *et al.* The Cohesin Release Factor WAPL Restricts Chromatin Loop Extension. *Cell* **169**, 693-707 e614, doi:10.1016/j.cell.2017.04.013 (2017).
69 Ryu, J.-K., Rah, S.-H., Janissen, R., Kerssemakers, J. W. J. & Dekker, C. Resolving the step size in condensin-driven DNA loop extrusion identifies ATP binding as the step-generating process. *bioRxiv*, 2020.2011.2004.368506, doi:10.1101/2020.11.04.368506 (2020).
70 Ainavarapu, S. R. *et al.* Contour length and refolding rate of a small protein controlled by engineered disulfide bonds. *Biophys J* **92**, 225-233, doi:10.1529/biophysj.106.091561 (2007).
71 Kuhn, W. Über die Gestalt fadenförmiger Moleküle in Lösungen. *Kolloid-Zeitschrift* **68**, 2-15, doi:10.1007/BF01451681 (1934).
72 Li, Y. *et al.* Structural basis for Scc3-dependent cohesin recruitment to chromatin. *Elife* **7**, 352, doi:10.7554/eLife.38356 (2018).
73 Nomidis, S. K., Carlon, E., Gruber, S. & Marko, J. F. DNA tension-modulated translocation and loop extrusion by SMC complexes revealed by molecular dynamics simulations. *Nucleic Acids Res* **50**, 4974-4987, doi:10.1093/nar/gkac268 (2022).

**Supplementary Video 1. Cohesin-mediated DNA loop extrusion blocked upon encounter of CTCF from its N-terminal side.** The example is the same as shown in Figure 2c. Raw image series and resulting kymographs of DNA and CTCF are shown. The images were median filtered with a 5 pixel window. The loop size as well as loop and CTCF positions were quantified as described in Material and Methods. The left cartoon illustrates the position of CTCF (the tip of the arrow represents its N-terminal side), the loop and the size of the loop. Note that the proportions are only roughly to scale and should be viewed as an illustration, rather than as an absolute quantification. The top view mimics the side-view cartoon and illustrates how changes in loop size are visible as changes in the intensity of the fluorescent ‘dot’ moving along DNA. Instead of varying brightness, loop size variations as represented as varying diameters of the loop ‘dot’.

**Supplementary Video 2. Cohesin-mediated DNA loop extrusion can pass CTCF.** When approached at ~50s, CTCF first blocks the extruding cohesin. Cohesin then retracts while its loop slightly shrinks and attempts again to pass CTCF at 66 s. This attempt was successful as CTCF was included in the loop and translocates with it.

**Supplementary Video 3. Repeated short CTCF-loop co-localization events.** Cohesin approaches CTCF and encounters it at 74 s, which blocks loop extrusion. Upon dissociation at 82 s, cohesin attempts to pass CTCF again at 105 s, 139 s, and 165 s. The encounter at 175 s results in passing of cohesin and CTCF translocates with the extruded loop.

**Supplementary Video 4. Stalling of loop extrusion at CTCF results in a direction switch of loop extrusion.** Upon encounter of cohesin with CTCF at 28 s, the loop continues to grow and the co-localized position of loop and CTCF moves downwards, away from the CTCF-cohesin encounter position. Cohesin and CTCF dissociate at 49 s.

**Supplementary Video 5. CTCF-stalled loops shrink in size upon dissociating from CTCF.** A previously extruded loop is shown. At 30 s, growth of the loop resumes and the loop encounters CTCF shortly after. The loop size and position remain constant while co-localized with CTCF. At 64 s, cohesin and CTCF dissociate and the loop shrinks after which it remains static.

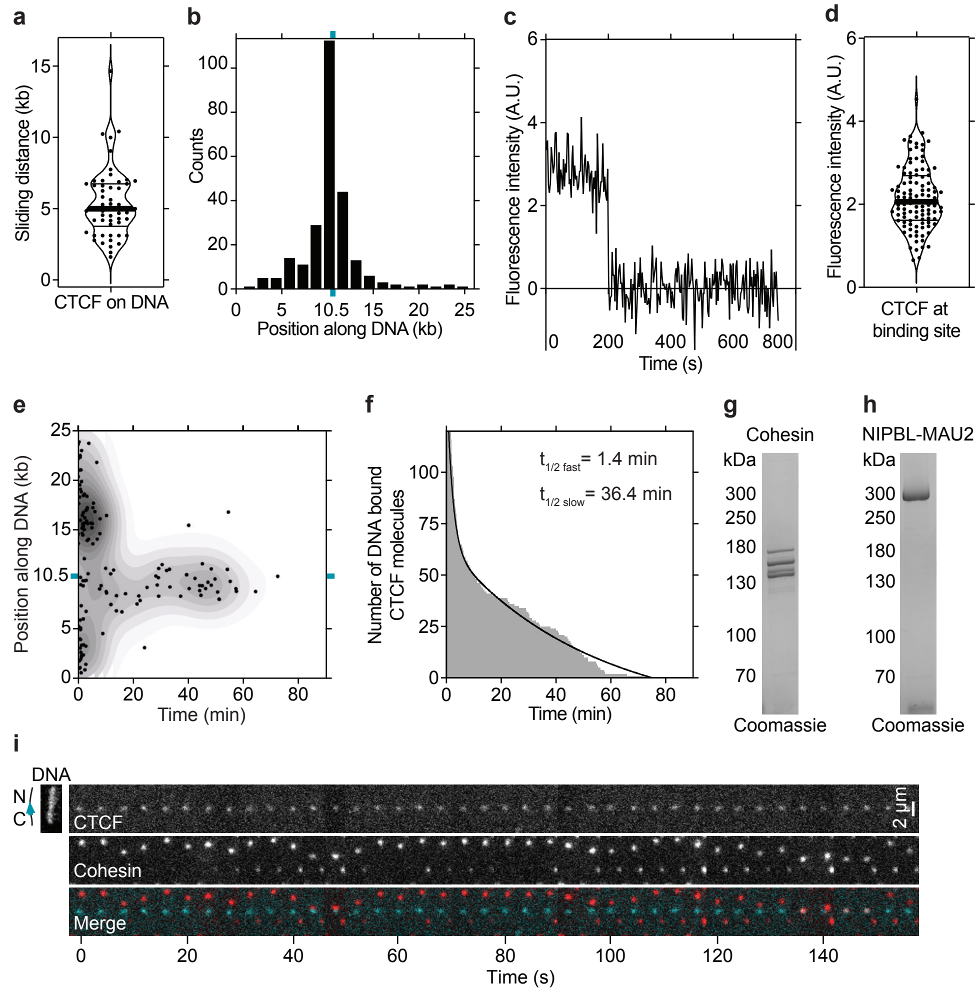


**Extended Data Figure 1. Recombinant CTCF characterization. a,** Distance (kb) travelled by TMR labelled CTCF molecules while diffusing before encountering the CTCF binding site. N = 54. **b,** Position of DNA bound TMR labelled CTCF following a brief wash step. The CTCF binding site (cyan tick) is at position 10,452 bp out of 26,123 bp. N = 251. The orientation of the DNA was determined using end-labelling by TetR as shown in Extended Data Figure 3f. **c,** Time trace of Alexa 660 (A660)-labeled CTCF signal bound at its DNA binding site bleaching in one step. **d,** Fluorescence intensity of A660 labelled CTCF signals at the CTCF binding site. N = 104. **e,** Residence time of TMR labelled CTCF on DNA. The CTCF binding site (cyan tick) is at position 10,452 bp out of 26,123 bp. N = 140. **f,** Residence time of TMR-labeled CTCF on DNA from (e) plotted as a histogram. Bi-exponential decay curve was fitted using Prism. **g - h,** Coomassie staining of recombinant cohesin and NIPBL-MAU2 after SDS-PAGE. **i** Example of cohesin diffusion blocked by CTCF. Cohesin and CTCF were labelled with A660 and TMR, respectively. Sytox Green DNA stain was introduced into the flow cell at the end of the experiment. This data is identical to main Figure 1f except it is formatted as a montage rather than as a kymograph.

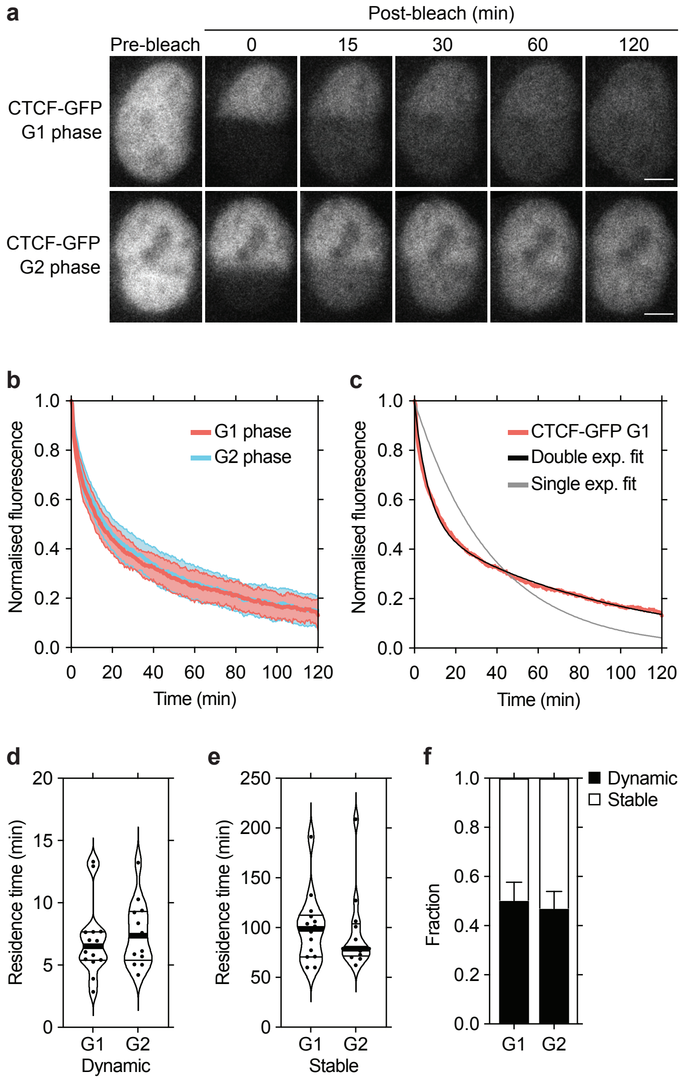


**Extended Data Figure 2. Chromatin-bound residence time of CTCF in vivo. a,** Images of inverse fluorescence recovery after photobleaching (iFRAP) in G1 and G2 phase CTCF-GFP HeLa cells. Scale bar, 10 µm. Half of the nuclear CTCF-GFP fluorescent signal was photobleached and the mean fluorescence in the unbleached and bleached regions was monitored by time-lapse microscopy. **b,** Graph showing the mean normalized difference in fluorescence intensity between the bleached and unbleached regions from cells treated as described in (a). Shaded regions represent standard error of the mean (s.e.m.). N ≥ 13 cells per condition. **c,** Graph showing single- and bi-exponential fits to G1 phase CTCF-GFP iFRAP data from (b)**. d - e,** Quantification of the residence time of (d) dynamically and (e) stably chromatin bound CTCF-GFP in G1 phase and G2 phase cells. **f,** Fraction of dynamically and stably bound CTCF-GFP in G1 phase and G2 phase.

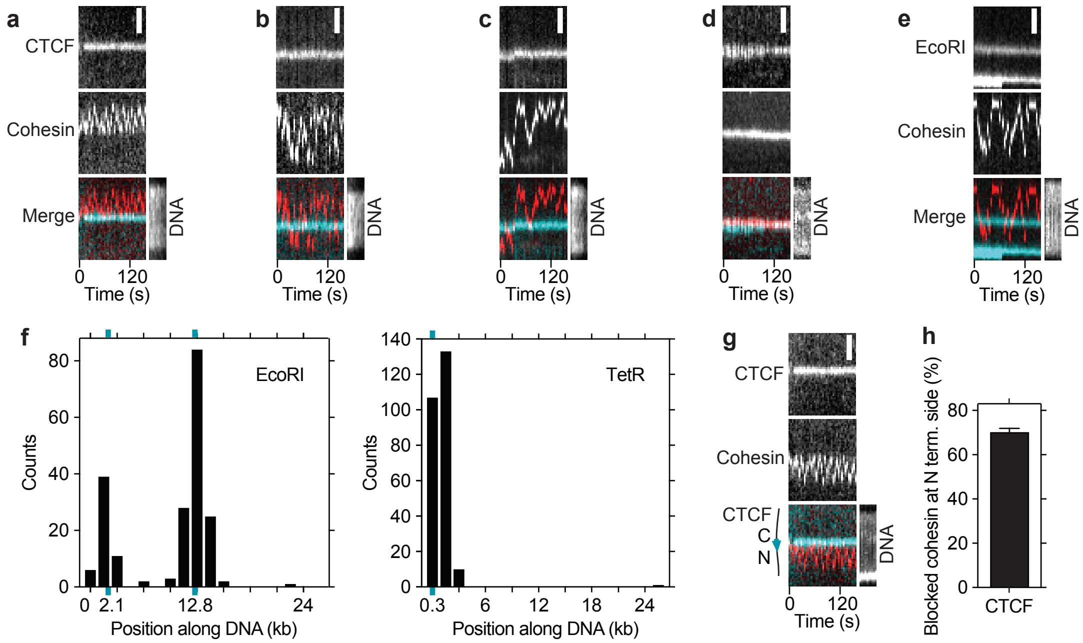


**Extended Data Figure 3. Cohesin diffusion assay characterization. a - d,** Examples of cohesin diffusion on DNAs with CTCF bound at its binding site. Cohesin was labelled with Alexa 660 (red). CTCF was labeled with tetramethylrhodamine (TMR) (cyan). Sytox Green DNA stain was introduced into the flow cell at the end of the experiment. Scale bar, 2 μm. (a) Example of cohesin diffusion blocked by CTCF. (b) Example of cohesin diffusing past CTCF multiple times within the imaging timeframe. (c) Example of cohesin diffusing past CTCF in one direction only. Example of cohesin diffusing past CTCF in one direction only. This behavior was observed very infrequently (2 ± 3 % of N = 264 events). This could be because cohesin-CTCF encounters were recorded after the system has reached equilibrium and so all the single-pass events had occurred before we could image them. If so, is unknown why some cohesin molecules behaved differently i.e. were able to pass CTCF multiple times (Extended Data Fig. 3B). (d) Example of cohesin-CTCF colocalization. **e,** Example of cohesin diffusing past TMR-labeled EcoRI$^{E111Q}$. **f,** Positions of DNA bound (left) Janelia Fluor 646-labeled EcoRI$^{E111Q}$ and (right) TMR-labeled TetR, which were flowed into flow cells at the end of diffusion experiments to determine the DNA orientation and hence the orientation of the CTCF binding site at position 10,452 bp. EcoRI restriction sites were present at positions 2,177 bp and 12,802 bp out of 26,123 bp. N = 201. Six TetO sequences were present at positions 40-274 bp. N = 251. **g,** As in main Figure 1f, except using a DNA in which the CTCF site was inverted. **h,** Fraction of blocked events that diffused on the DNA between the tether point and the N terminal side of CTCF using the DNA template as used in (g) (mean ± SD from 3 (N=48) independent experiments).

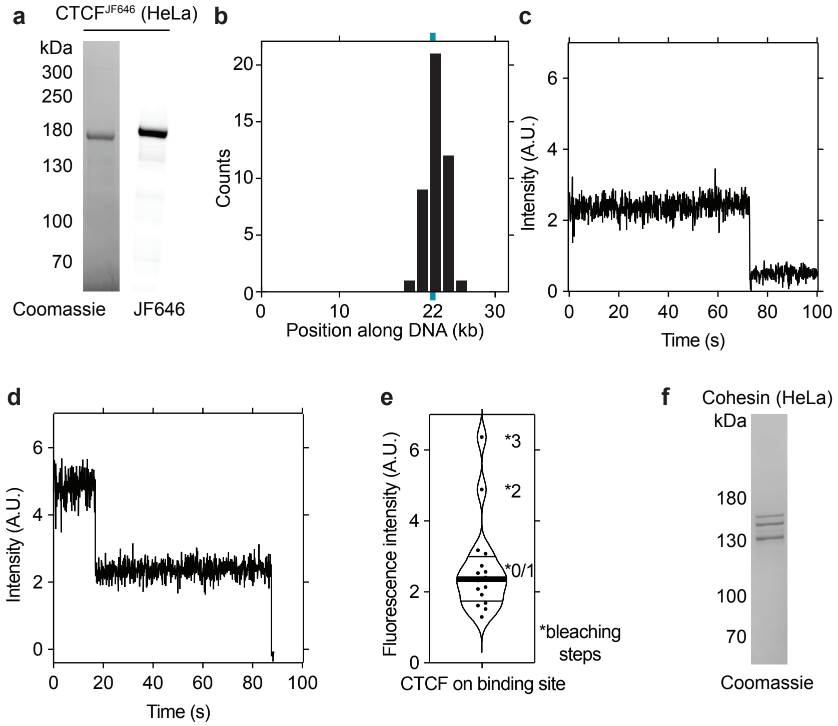


**Extended Data Figure 4. HeLa CTCF characterization. a,** Coomassie staining of HeLa CTCF after SDS-PAGE. JF646 was visualized by epi-red excitation. **b,** Position of DNA bound JF646-labeled CTCF following a wash step with a buffer supplemented with 220 nM Sytox Orange. The CTCF binding site (cyan tick) is at position 9,667 bp out of 31,767 bp. N = 251. **c,** Time trace of JF646-labeled CTCF signal bound at its DNA binding site bleaching in one step. **d,** Time trace of JF646-labeled CTCF signal bound at its DNA binding site bleaching in two steps. **e,** Fluorescence intensity of JF646-labeled CTCF signals at the CTCF binding site. N = 16. **f,** Coomassie staining of HeLa cohesin after SDS-PAGE.

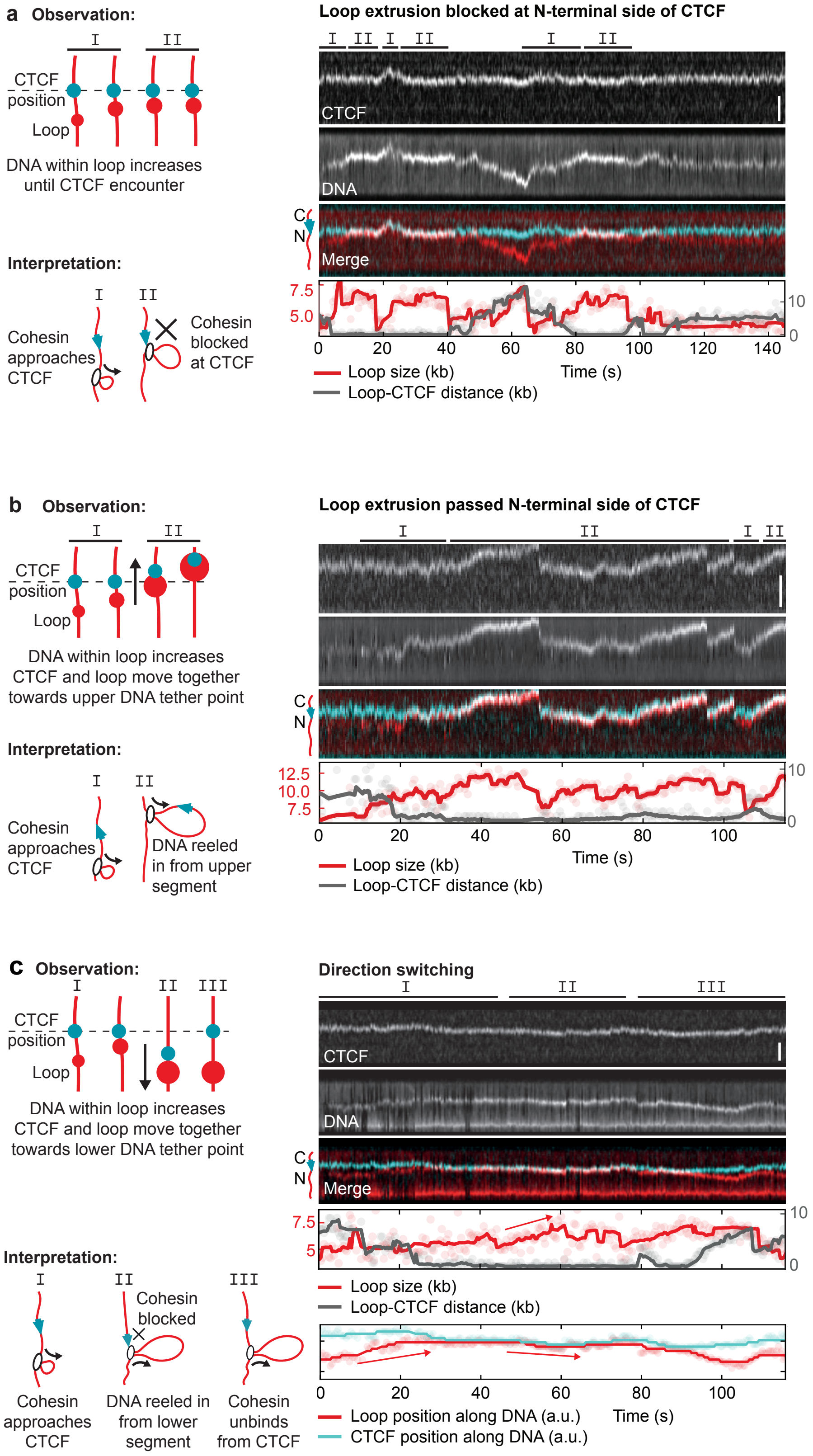


**Extended Data Figure 5. Additional examples of loop extrusion blocking, passing and direction switching upon encountering CTCF. a - c,** (Left panels) observation and interpretation illustrations of (right panels) kymographs of cohesin-mediated DNA loop extrusion encountering N-terminally oriented CTCF (cyan) labeled with Janelia Fluor 646 (JF646). DNA loops were visualized by Sytox Orange stain. Scale bar, 2 µm. (a) Growth of the DNA loop stops upon encountering CTCF at timepoints ~ 12 – 18 s, 22 – 40 s and 82 – 95 s. (b) The DNA loop continues to grow upon encountering CTCF at 31 s, and CTCF passes into the loop and translocates with it. (c) The growing loop encounters CTCF at 28 s. CTCF and the growing DNA loop move towards the lower DNA tether point, indicating extrusion on the side facing away from CTCF.

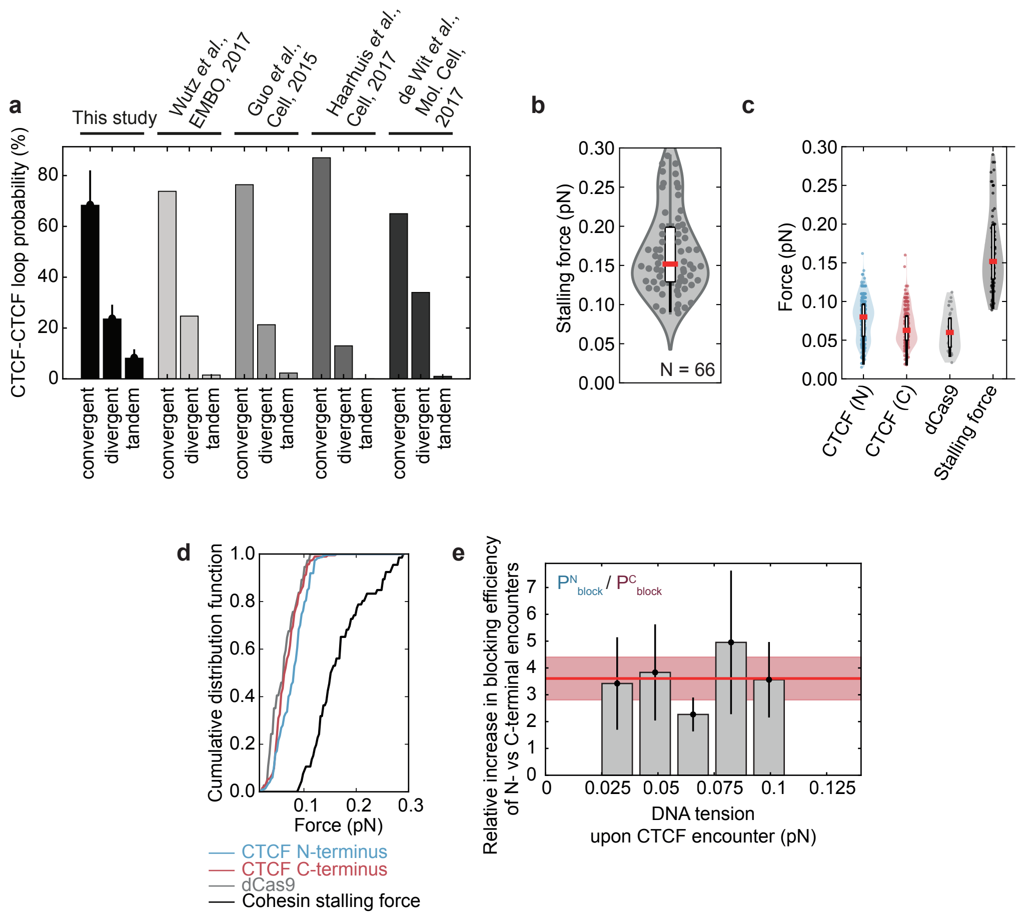


**Extended Data Figure 6. Stalling force of cohesin and force sampling for encounters with the N-/C-terminus of CTCF and dCas9. a,** Combinatorial loop extrusion blocking efficiency at a pair of CTCF sites oriented in a convergent (><), tandem (>> and <<), and divergent (<>) manner. The percentages were obtained by multiplying the blocking probability of N- and C-terminal encounters in the force range 0.04-0.08 pN, as depicted in Figure 2E, and normalizing to 100% (see Methods). The relative fraction of CTCF-anchored loops in that we obtained from the single-molecule experiments are compared to published values extracted from Hi-C data[4, 66-68]. **b,** Stalling force of cohesin. Red mark denotes the median value and whiskers extend to the first and third quartiles. **c,** The DNA tension measured at encounters of loop-extruding cohesin with the N- and C-terminus of CTCF and dCas9. The stalling force values from panel B is shown as well for comparison. N = 297, 184, 37, 66 for CTCF (N), CTCF (C), dCas9 and the stalling force measurements, respectively. **d,** The empirical cumulative distribution function (eCDF) of the data shown in panel C. **e,** Ratio of the N-terminal and C-terminal blocking probabilities. N-terminal encounters block loop extrusion 3.6 ± 0.8 -fold (mean ± SD) more often than encounters from CTCF's C-terminal side, independently of DNA tension.

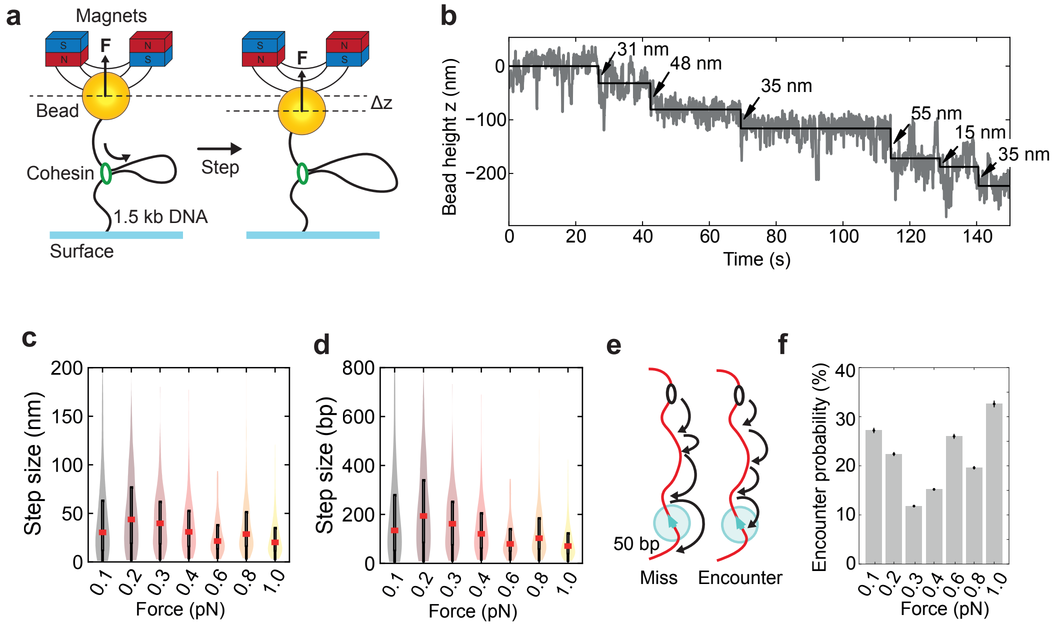


**Extended Data Figure 7. The force-dependent step size of cohesin loop extrusion cannot solely explain the observed force dependence of CTCF blocking loop extrusion. a,** Magnetic tweezers setup to observe individual loop extrusion steps by human cohesin, depending on the applied force, based on *(68)*. The change in bead height Δz corresponds to steps by cohesin. **b,** Example magnet tweezer trace showing stepwise changes in bead height in the presence of cohesin, NIPBL-MAU2 and ATP. Line denotes steps fitted using the step-finding algorithm. **c,** Step sizes in nanometers as measured by Magnetic Tweezer experiments, for various applied forces ranging from 0.1 pN to 1 pN. **d,** Step sizes versus force from (c), but converted to base pairs. **e,** Simulation setup: starting from a randomly chosen binding position along DNA, cohesin takes steps along DNA, which are sampled from the measured step size distribution. An 'encounter' is considered if cohesin comes within 50 bp of CTCF. Under the lenient assumption that the CTCF N-terminus is unstructured and may be approximated by a freely jointed chain, its radius of gyration $R_G$ is estimated using the $N_K$ = 268 amino acids from the N-terminus to zinc finger 1[26], with a contour length of $l_K$ ~0.4 nm per amino acid[70], resulting in $RG = N_k l_K^2/6$ ~7 nm[71]. This distance corresponds to roughly 20 bp, given the contour length of a basepair of 0.3 nm. A threshold of 50 bp was thus conservatively chosen because the CTCF N-terminus may be as long as 14 nm but is likely more compact due to folding of the CTCF N-terminus. The simulations thus likely represent an upper limit of the encounter probability. **f,** Simulated encounter probability of cohesin and CTCF. Note that the encounter probability does not exceed ~40%, even at the smallest step size distribution (measured at 1 pN). In contrast, the blocking probability of N-terminal encounters of cohesin and CTCF increases from 0 to 100% within 0-0.14 pN (Figure 2g). Force-dependent step sizes of cohesin can thus not solely explain the observed N-terminal blocking probability. We therefore suspect that DNA tension increases the blocking efficacy of CTCF by other mechanisms, such as by reducing not only cohesin's step size but also the frequency with which it takes steps, thus providing more time for CTCF and cohesin to bind to each other; or by reducing thermal fluctuations of DNA[48], which could reduce the space that CTCF has to explore to find cohesin. It is also conceivable that cohesin's 'motor' activity can overcome the low 1 µM binding affinity of CTCF-cohesin interactions[72] more easily at low DNA tension than at high tensions, which are close to the stalling force of loop extrusion, and at which cohesin has to generate higher forces to extrude DNA. Finally, DNA tension could also change cohesin's responsiveness to CTCF by influencing how cohesin performs loop extrusion[73].

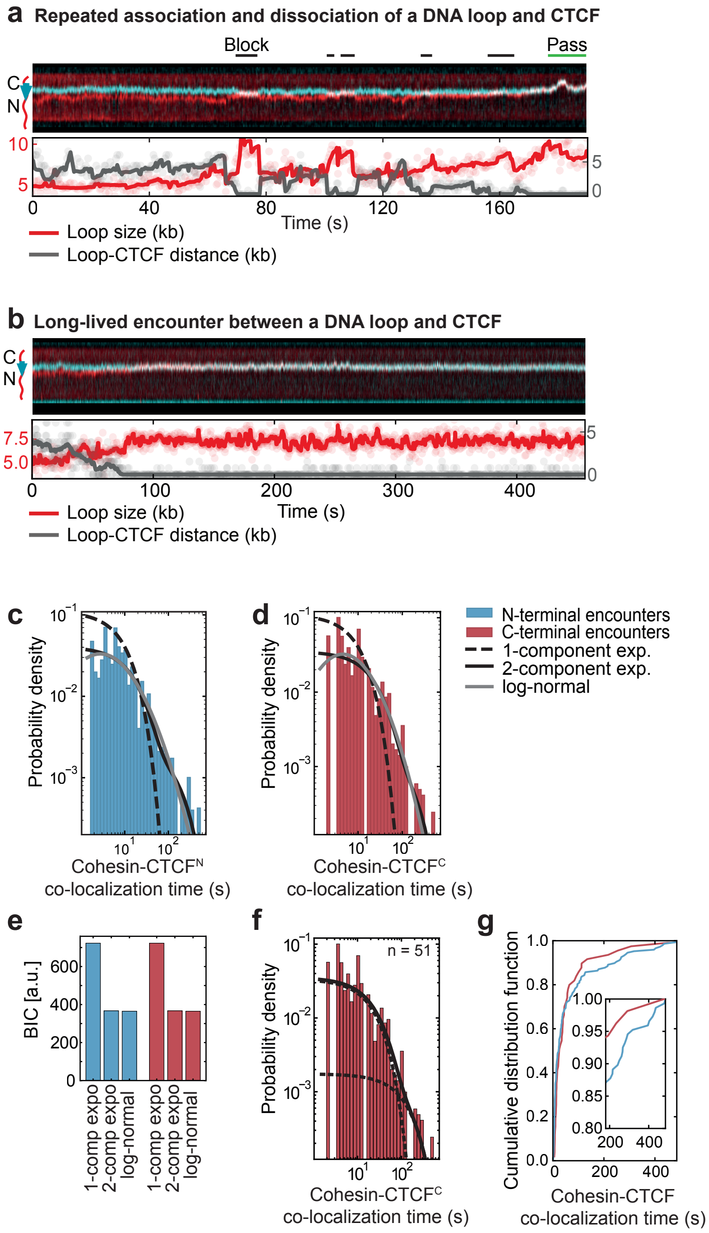


**Extended Data Figure 8. Cohesin-CTCF residence time characterization. a,** Example of repeated approaching of CTCF (cyan) by cohesin, blocking of further loop extrusion and dissociation of the cohesin-CTCF interaction. Cohesin passes CTCF at the end of the kymograph. DNA loops (red) were visualized by Sytox Orange stain. Scale bar, 2 µm. **b,** Example of a growing loop encountering CTCF (cyan), stalling and co-localizing until the end of image acquisition. DNA loops (red) were visualized by Sytox Orange stain. Scale bar, 2 µm. **c,** Co-localization times of cohesin for the encounters from the N- and **d,** C-terminal side of CTCF (N = 147 and N = 51 for N- and C-terminal encounters, respectively). The distributions are fitted to a mono-exponential, bi-exponential and log-normal distribution. **e,** Bayesian Information Criterion (BIC) for the three models on N- and C-terminal encounters. Notably, both a 2-exponential as well as a lognormal distribution fit the distributions equally well. The parameters of the lognormal fits of the form $(x\sigma\sqrt{2\pi})^{-1}\exp((\ln(x)-\mu)^2/(2\sigma^2))$ are μ=1.472 s,σ=3.181 s for N-terminal and μ=1.246 s,σ=3.066 s for C-terminal encounters. **f,** The residence time of encounters between cohesin and CTCF's C-terminus is well described by a bi-exponential distribution with rate constants k1 = 0.043 s-1 and k2 = 0.008 s-1 (τ1 = 23 s and τ2 = 125 s). **g,** Cumulative distribution function of the cohesin-CTCF co-localization time for N- (blue) and C-terminal (red) encounters. Inset: magnified view of co-localization times ≥3 min.

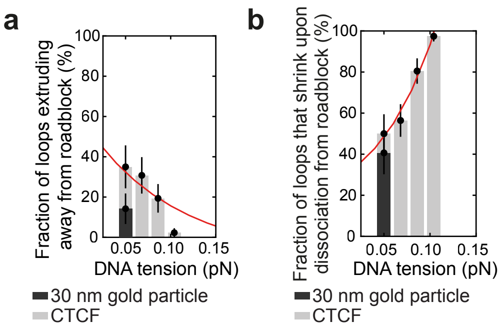


**Extended Data Figure 9. Frequency of direction switching and loop shrinkage following encounters between cohesin and CTCF or gold nanoparticles. a,** The fraction of loops extruding on the side facing away from CTCF (grey bars) or 30 nm gold nanoparticles (black bar; 14±8 % [mean ± SD]). CTCF data is replotted from Fig. 3C. Encounters with gold nanoparticles over a force range of 0.02-0.05 pN were reanalyzed from[45] (N = 21). **b,** The fraction of loops which shrink upon release from CTCF (grey bars) or 30 nm gold nanoparticles (black bar; 41±10 % [mean ± SD]) versus DNA tension at the moment of encounter. CTCF data is replotted from Fig. 3E. Encounters with gold nanoparticles over a force range of 0.02-0.05 pN were reanalyzed from[45] (N = 22).

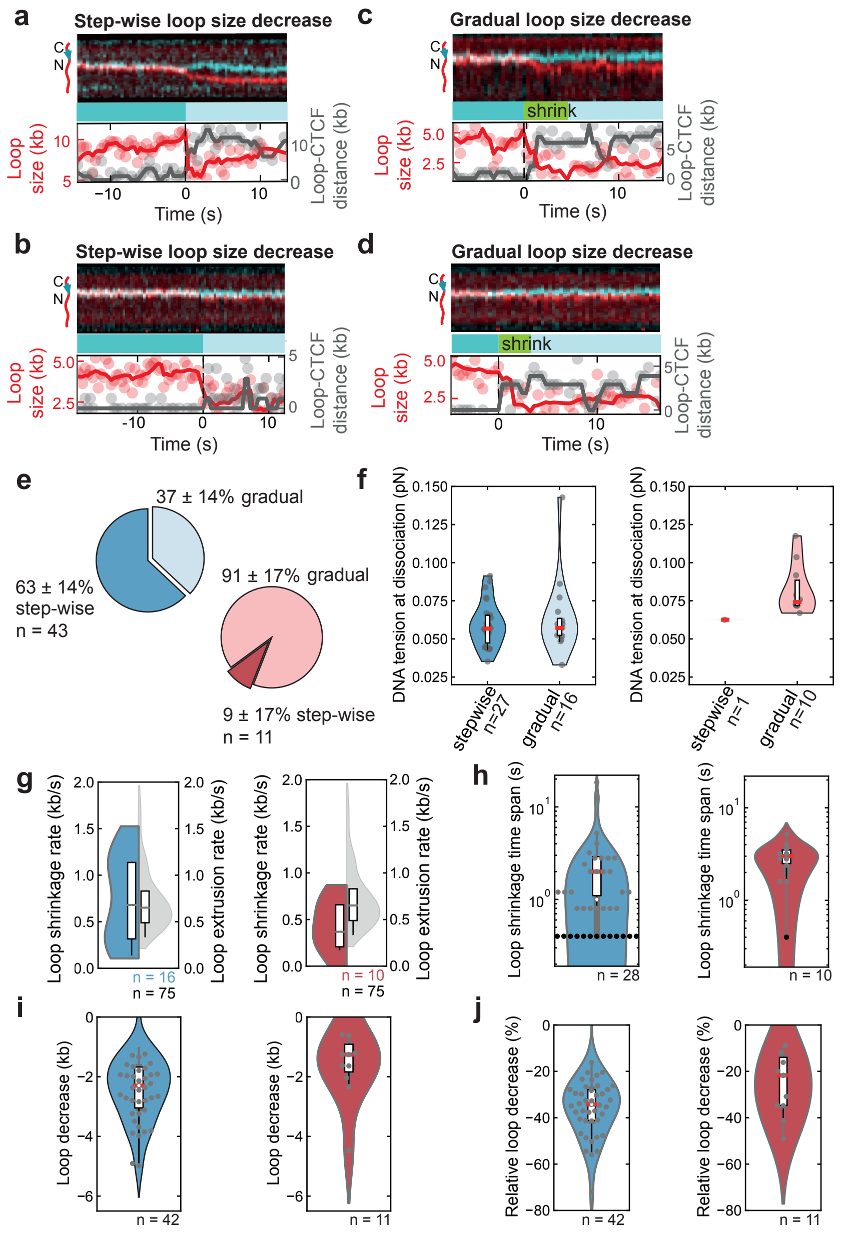


**Extended Data Figure 10. Kinetics of cohesin-CTCF dissociation. a - b,** Examples of step-wise and **c - d,** continuous loop shrinkage upon dissociation of cohesin from CTCF. Scale bar, 2 µm. Related to Figure 3b, e. **e,** The fraction of step-wise and continuous loop shrinkage for encounters from the N-terminal (blue) and C-terminal (red) side. **f,** DNA tension for loops which shrink step-wise or gradually. There is no statistically significant difference in DNA tension between the two modes (p>0.05, 2-sample Kolmogorov–Smirnov test). **g,** Loop shrinkage rate, in comparison to cohesin loop extrusion rate (grey), and **h,** distribution of shrinkage time spans. Black dots represent step-wise shrinkage events that happen within one imaging time interval, i.e. 0.4 s. **i,** Absolute and **j,** relative loop size decrease for N- and C-terminal encounters in blue and red, respectively.
